# Quantification of lipid sorting during clathrin-mediated endocytosis

**DOI:** 10.1101/2025.03.04.641423

**Authors:** H. Mathilda Lennartz, Susana da Costa Nunes, Kristin Böhlig, Sascha M. Kuhn, Lior Moneta, Wan Yee Yau, Lukáš Opálka, Juan M. Iglesias-Artola, Carl D. Modes, Alf Honigmann, André Nadler

## Abstract

Clathrin-mediated endocytosis is a major transport route for proteins from the plasma membrane to the interior of the cell. While the recruitment of cargo proteins to clathrin-coated pits is well understood, it remains an open question if lipids are also sorted by this process. To address this question, we combined super-resolution STED imaging of bifunctional lipid probes with mathematical modeling. Quantification of 10 different lipid species revealed significant differences in pit partitioning, ranging from slight enrichment to moderate exclusion. We find that the lipid asymmetry in the plasma membrane is sufficient to explain the observed trend. Taken together, our findings imply that clathrin-mediated endocytosis has a minor selectivity for cytoplasmic leaflet lipids, but overall does not significantly contribute to lipid sorting compared to non-vesicular trafficking. More broadly, we believe that lipid super-resolution imaging will be a powerful approach to quantify lipid partitioning into membrane structures in cells.

## Introduction

Clathrin-mediated endocytosis (CME) is the primary route of bulk cellular ingestion and lipid uptake during retrograde vesicular transport. More than 50 proteins are involved in the formation of clathrin-coated vesicles with a diameter of 60-120 nm. They are recruited to the endocytic site in a highly controlled temporal order during the three stages of initiation, maturation, and fission ^1–4^.

Cargo proteins are enriched in clathrin-coated pits (CCPs) by specific interactions with adaptors via short peptide motifs and/or posttranslational modifications ^5,6^. Much less is known about the extent and the mechanisms of lipid sorting during CME. Previous studies have shown that PI(4,5)P2 localization to CCPs is a necessary precondition for pit initiation and maturation. However, there is no compelling evidence for PI(4,5)P2 enrichment in the pit ^7–9^. Further reports suggest that lipids associated with ordered membrane domains are depleted in CCPs, such as cholesterol, gangliosides, and glycosylphosphatidylinositol, whereas lipids associated with disordered membrane domains should accumulate ^10–14^. These studies are based on indirect evidence for lipid sorting into CCPs by monitoring the endosomal levels of fluorescent tracers such as glycophosphatidylinositol-anchored proteins, fluorescent lipid analogs, and lipid-binding toxins. Notably, reporters such as cholera toxin cluster lipids, which affect sorting and uptake mechanisms ^15^. Fluorescent lipid analogs do not reliably reflect the behavior of native lipids, as the introduction of a bulky chromophore alters the chemical properties of the parent molecule ^16–18^. More generally, quantification of lipid uptake on the level of endosomal structures convolutes plasma membrane and endosomal sorting mechanisms. A systematic investigation of lipid sorting during CME based on direct evidence at the pit has yet to be carried out for a broad selection of structurally diverse lipid species.

Here, we use our recently introduced approach for imaging cellular lipid localization ^19,20^ based on bifunctional lipid probes ^21–23^ to directly quantify lipid partitioning into CCPs. We adapted our methodology for super-resolution stimulated emission depletion (STED) microscopy and captured temporally resolved CCP maturation for a library of 10 different lipid species to investigate lipid enrichment in early and late-stage CCPs. Combination with theoretical modeling allowed us to quantify true lipid enrichment from the apparent enrichment caused by microscope convolution of the membrane geometry during pit maturation. Our analysis revealed significant differences between lipid species, suggesting differential partitioning into CCPs. By comparing different sorting mechanisms, we conclude that differential lipid partitioning into CCPs is primarily driven by the lipid asymmetry of the plasma membrane. Lipid partitioning into CCPs is thus fully determined by plasma membrane lipid composition and transbilayer lipid distribution, implying that CME combines selective protein and non-selective lipid transport.

## Results

### Super-resolution STED resolves lipid species localization in clathrin-coated pits

To quantify the role of CME for sorting lipids at the plasma membrane, we optimized our recently developed lipid imaging workflow ^20^ for super-resolution STED microscopy. Briefly, near-native bifunctional lipid species modified with diazirine and alkyne moieties were loaded to the outer leaflet of the plasma membrane and fixed via subsequent UV-light-and chemical crosslinking (Figure 1A). To restrict lipid localization to the plasma membrane, lipids were UV crosslinked shortly after lipid loading, and cells were chemically fixed according to Mund and Tschanz et al. ^24^ to ensure conservation of CCP ultra-structure. UV-crosslinking time was minimized using high-powered LEDs at 365 nm with a collimation lens to maximize photon flux (Extended Data 1, Supplementary Data 1). Subsequently, samples were permeabilized to remove non-crosslinked probes and to allow for immunolabelling of CCP reporter proteins. Lipid-protein conjugates were fluorescently labeled by repeated copper-catalyzed click chemistry (in the following referred to as click chemistry) (Extended Data 2, 3). UV-crosslinking and fixative addition were completed within < 10 seconds, which allowed to capture snapshots of CCPs at different maturation stages ^25^.

We used the well-characterized SK-MEL2 dynamin-2 GFP reporter cell line ^26^ in combination with immunostaining of clathrin and AP2 to classify CCPs according to their maturation stages. Structures positive for AP2 and clathrin were classified as early-stage pits ^25,27^, whereas structures additionally positive for dynamin-2 were classified as late-stage pits ^25,28^. Structures positive only for clathrin were classified as clathrin-only structures (Figure 1B, C). Lipids and clathrin were imaged at super-resolution (STED), while dynamin-2 and AP2 were acquired in diffraction-limited confocal mode (Extended Data 4). Taken together, this workflow allowed us to resolve lipid species in CCPs in different maturation stages.

**Figure 1:**
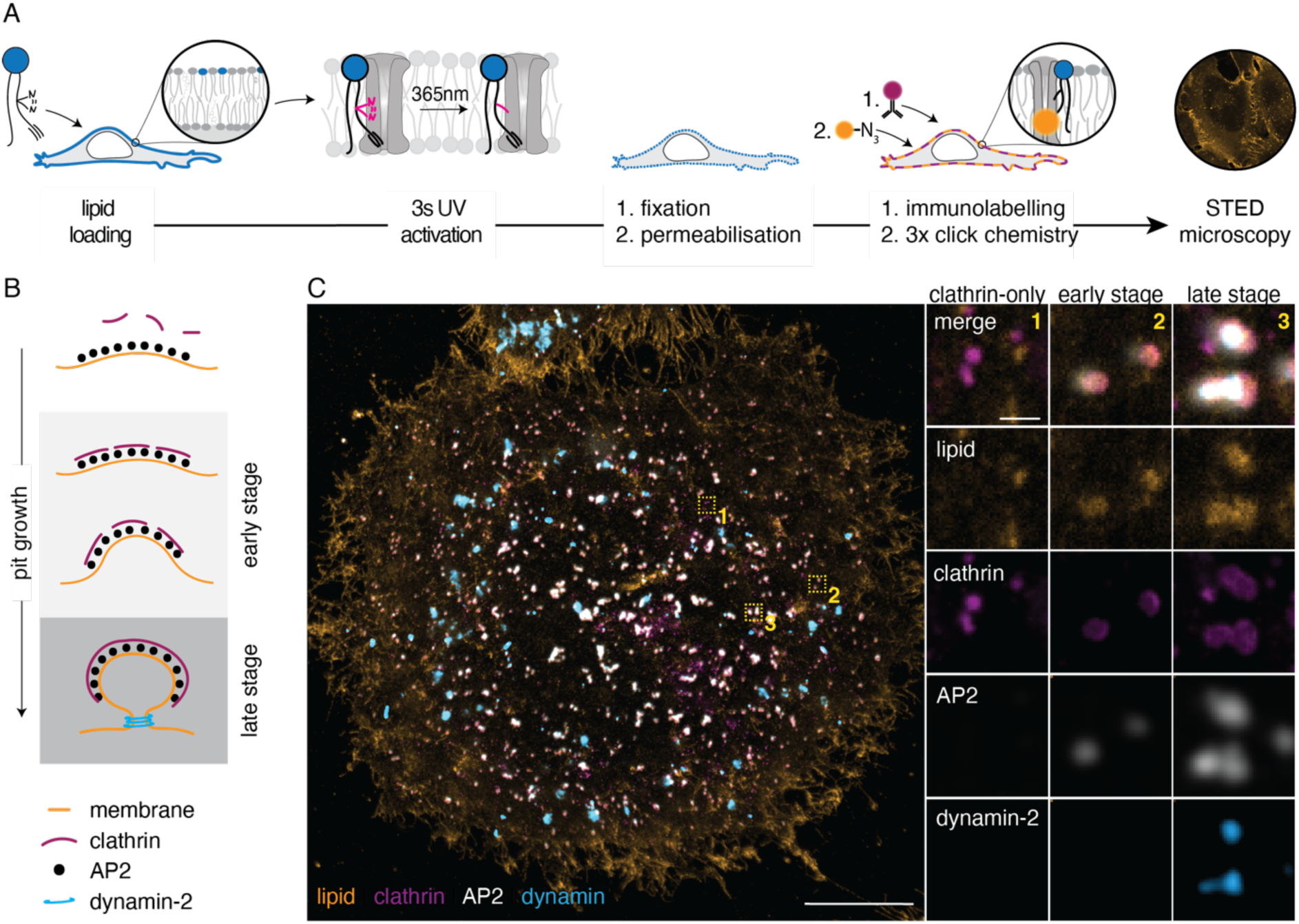
Super-resolution lipid imaging of lipid localization in clathrin-coated pits. A: Super-resolution lipid imaging workflow. Lipid probes were loaded into the plasma membrane of living cells using α-methylcyclodextrin. Photo-crosslinking to proteins with a 3s UV light pulse was followed immediately by chemical fixation. Samples were permeabilized, stained by immunolabelling and click chemistry, and imaged using 4-color confocal microscopy with the lipid and clathrin-channel super-resolved by STED. B: Stages of the growth of a CCP were classified using protein markers into clathrin-only structures, early-stage CCPs positive for AP2 and clathrin, and late-stage CCPs positive for AP2, clathrin, and dynamin-2. C: A representative STED image of the basal membrane of a SK-MEL2 cell loaded with bifunctional PC(16:0|Y) (orange) is shown, stably expressing dynamin-2-GFP (blue). Clathrin (purple) and AP2 (grey) were immunolabelled. Scale bars: 10µm (full image), 500nm (magnified areas).

### Quantification of lipid partitioning into clathrin-coated pits

To quantify lipid species partitioning into CCPs, we next segmented the highly resolved clathrin signal. We classified each pit based on the co-localization of the clathrin mask with the respective AP2 and dynamin-2 signal into clathrin-only structures, early (AP2 and clathrin positive) and late (AP2, dynamin-2, and clathrin positive) CCPs (Figure 2A). Mean lipid signal intensities were determined for all pits over the segmented clathrin mask and the surrounding bulk plasma membrane in a square of 1.5 µm x 1.5 µm (ROI), excluding other pits and unidentified bright structures. Apparent lipid partitioning (P) was defined as the ratio of the mean lipid signal intensity within each clathrin mask and the surrounding plasma membrane. We found a general trend of increasing apparent lipid partitioning from early to late-stage pits (Extended Data 5, 6). This is shown using PC(16:0|Y) as an example, a probe closely mimicking a common plasma membrane PC species, with an apparent partitioning from P = 1.64 ± 0.45 (median ± quartile) to P = 2.09 ± 0.36 for early and late-stage pits, respectively (Figure 2B, C, G). Next, to delineate to which extent this effect was indicative of true lipid accumulation in CCPs or whether it was a reflection of the changing membrane geometry during pit maturation, we modeled the convolution of the membrane geometry by the PSF of our microscope (Figure 2D-F).

Geometries of early and late-stage pits were modeled by combining the shapes of partial orbs (pit dome) and catenoids (pit neck if applicable) using previously reported EM data of CCP membrane ultrastructures obtained from the SK-MEL2 cell line ^29^ (Figure 2F, G, Materials and Methods). Pits were classified as late-stage if their neck diameter measured less than 40 nm, a curvature that has been reported to facilitate dynamin membrane association *in vitro* ^30^. Next, an even distribution of point emitters per surface area was assumed across the 3D models, representing an even lipid distribution. From these distributions, STED images were simulated according to the PSF of our microscope. This model predicted a median (± quartile) of the apparent partitioning of 1.82 ± 0.51 for early stages and 2.10 ± 0.25 for late stages (Extended Data 7), similar to the experimentally acquired values of PC(16:0|Y). The modeled pit geometries and resulting apparent partitioning values are not normally distributed, presumably because the distribution directly reflects the original EM dataset, which contains low numbers of individual pit geometries during maturation.

A closer inspection of the modeled pit structures showed that late-stage pits are much more consistent in their geometry compared to the early stages (Figure 2F). The early-stage pits contain a mixture of two geometries (domes without necks and domes with necks), but do not capture any transition geometries. A similar effect was not observed for late-stage CCPs, which include pits of similar geometries, indicating a more faithful recapitulation of the late-stage pit structures by the model.

To convert the convoluted apparent experimental partitioning values into absolute partitioning values, we normalized the apparent partitioning distribution to the median of the modeled distribution (Figure 2G, Extended Data 8). For PC(16:0/Y) this normalization provided a median partitioning value close to one for early and late-stage pits (Figure 2G), indicating close to even partitioning of the lipid between the plasma membrane and the CCP.

Taken together, modeling pit geometry from known reference data allowed us to obtain absolute lipid partitioning values from convoluted experimental measurements, a key prerequisite for quantitatively studying the role of lipids in membrane compartmentalization.

**Figure 2:**
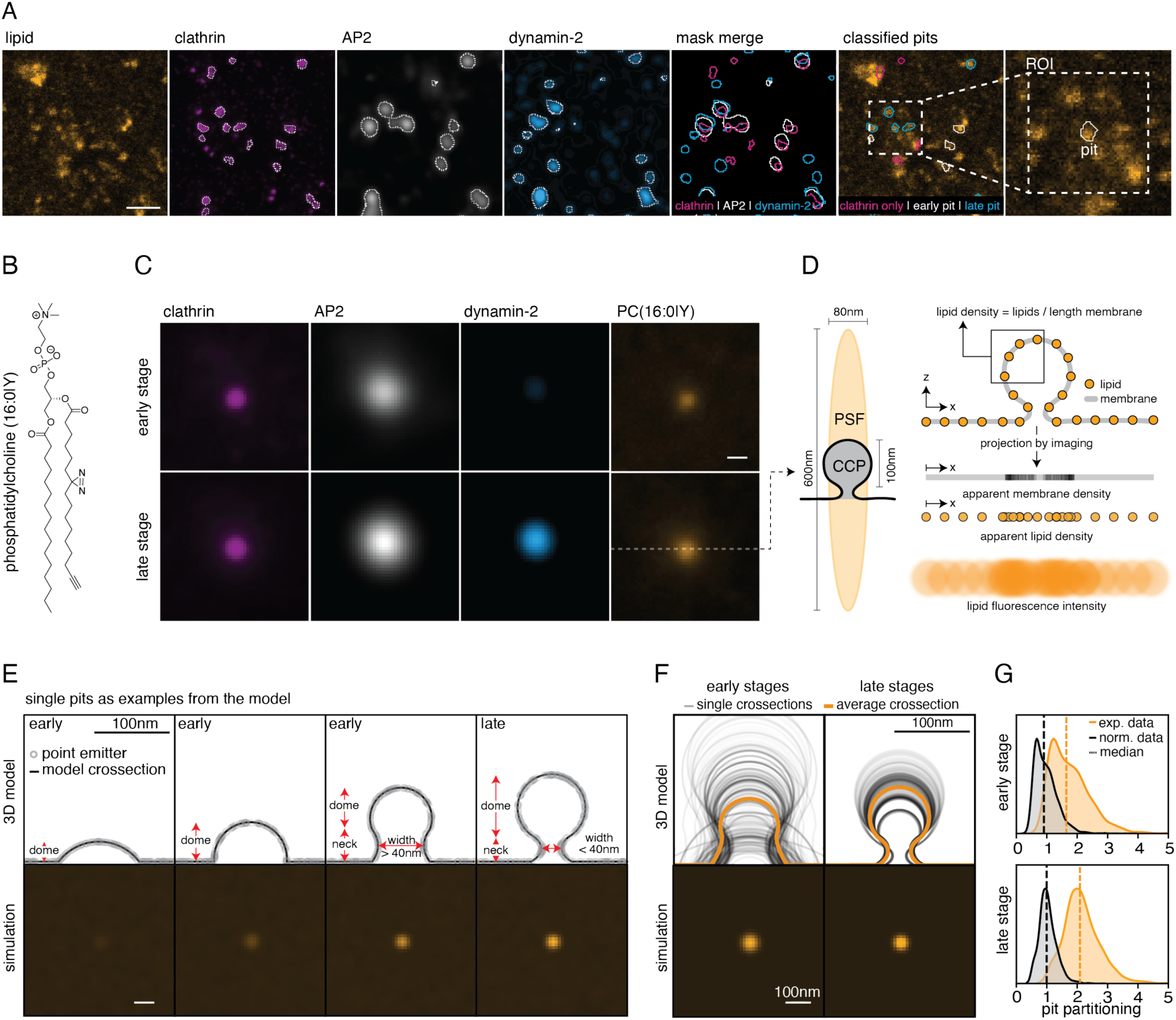
Quantification of lipid partitioning in clathrin-coated pits. A: Image analysis pipeline. The super-resolved clathrin signal (purple) was used to generate a segmentation mask (white dotted overlay). Pits were classified as clathrin only (purple), early-stage (white), or late-stage pits (blue), depending on their overlap with the AP2 and dynamin-2 masks generated from the respective channels. Apparent lipid partitioning was calculated by measuring the mean lipid intensities of individual pits and division by the mean lipid intensity of the surrounding plasma membrane (ROI). Scale bar: 1µm. B: Chemical structure of PC(16:0/Y). C: Average 4-color fluorescence images for early-and late-stage pits. Fluorescent images of CCPs were centered on the centroid of the clathrin mask before averaging. Scale bar: 100 nm. D: A scheme is shown on the left for the dimensions of the experimentally determined point spread function of the microscope in relation to a late-stage CCP. On the right, the process of convolution during imaging is depicted. A pit cross-section with even point emitter distribution is projected into one dimension, resulting in higher apparent membrane densities and lipid densities at the region of the pit invagination. After blurring by the PSF of the microscope, the pit appears brighter compared to the surrounding flat membrane. Lipid density as a readout for lipid partitioning must be determined as the ratio of lipids over surface area. E: Top panels: Cross-sections of modeled CCPs. Bottom panels: Simulated STED for the displayed pit geometries. Scale bars: 100 nm. F: Average cross-sections (black) of all simulated pits (grey) and averaged simulated images are shown. Scale bar 100 nm. G: The pit partitioning distributions of early and late-stage pits are shown for PC(16:0|Y) before (orange, exp. data) and after (grey, norm. data), normalizing to the median of the model. The respective median of the data is shown as a dotted line.

### Structurally diverse lipids partition differentially into clathrin-coated pits

To compare the partitioning of different lipid species into CCPs, we next studied a library of 10 bifunctional lipids that cover a large chemical space. The library included Glucosylceramide (GlcCer(Y)), sphingomyelin (SM(Y)), phosphatidylethanolamine (PE(18:1|Y)), a plasmalogen PC (pPC(18:1|Y)), as well as PC species of differential chain length, saturation degree and fatty acid positioning (Figure 3A, Materials and Methods) ^19,20^.

Focusing on the quantification of absolute lipid partitioning for late-stage pits we found that median partitioning values (± quantiles) ranged from 1.07 ± 0.16-fold to 0.84 ± 0.17-fold (Figure 3B). The magnitude of the observed partitioning of lipids into CCPs is also reflected in temporally highly resolved uptake kinetics of all lipids into early endosomes (Supplementary Figures 1, 2), which revealed no significant differences in uptake rates and only small differences in maximal endosomal lipid levels.

While the differences in pit partitioning are subtle, statistical analysis showed that most differences are highly significant (Figure 3C, Extended Data 9). Sorting lipid species by CCP partitioning provided the following order: PC(18:0|Y) > pPC(18:1|Y) > GlcCer(Y) > PC(20:0|Y) > PE(18:1|Y), PC(16:0|Y) > SM(Y) > PC(18:1|Y) > PC(Y|20:4) > PC(20:4|Y).

Overall, it appears that polyunsaturated lipids are slightly excluded from CCPs while saturated lipids tend to be slightly enriched in CCPs. However, this observation alone does not allow for the assignment of mechanistic roles of structural features such as headgroup, fatty acid saturation degree, or chain length for CCP partitioning. Taken together, quantification of lipid partitioning in late CPPs for 10 lipid species revealed significant differences of up to 21% between species, suggesting that lipids partition into CCPs in a species-specific manner.

**Figure 3:**
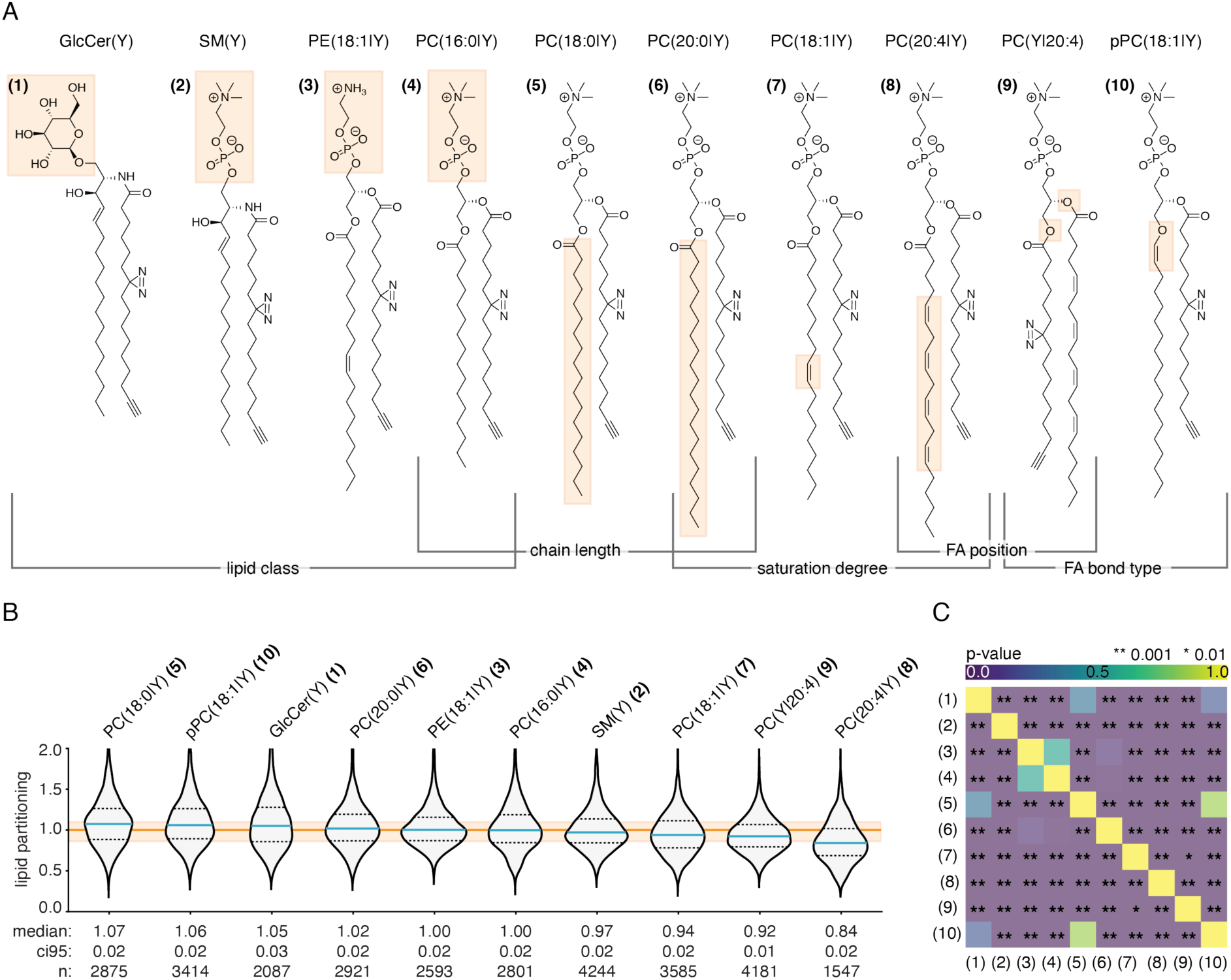
Comparison of 10 different lipid species reveals differential partitioning in clathrin-coated pits. A: Lipid library used to screen lipid enrichments in CCPs for structurally different lipids. Structural differences are highlighted in orange. Y – bifunctional fatty acid, GlcCer – glucosylceramide, SM – sphingomyelin, PE – phosphatidylethanolamine, PC – phosphatidylcholine, pPC – plasmalogen phosphatidylcholine. B: Violin plots of normalized late-stage CCP lipid partitioning distributions for all lipid probes. Distributions are ordered from highest to lowest median partitioning. Median values (blue line) with the respective quartiles (dotted line) are given below the plots with the 95% confidence intervals (ci95). The normalized median of the modeled distribution for a generic lipid evenly distributed between the plasma membrane and CCP is indicated by the orange line, with its respective quartiles (shaded area). C: Statistical significance between all lipids was determined by a two-sided randomized permutation test. P-values are color-coded, ** p-value < 0.001, * p-value < 0.01. Lipid species are indicated by numbers, as defined in A.

### Lipid partitioning correlates with lipid melting temperature and lipid asymmetry

To understand whether general features could be responsible for the observed differences in CCP partitioning, we performed a correlation analysis with known biophysical properties of the analyzed lipid species. We considered three possible mechanisms of sorting: partitioning of lipids into curved membranes driven by lipid shape (lipid resting curvature), sorting due to phase separation in curved membranes (lipid melting temperature), and sorting due to the asymmetric trans-bilayer distribution of lipids in the plasma membrane (lipid asymmetry) (Figure 4A, B). Values of lipid resting curvature, lipid melting temperature, and trans-bilayer lipid distribution were obtained from the literature ^31–36^ (Supplementary Data 2 - 4). In cases of missing reference data, values were approximated based on the structurally closest lipid counterparts.

First, we considered sorting by lipid shape. According to this hypothesis, cone-shaped (e.g., lyso-phosphatidic acid) and inverse cone-shaped lipids (e.g., 1,2-dioleoyl-sn-glycero-3-phosphoethanolamine) would be sorted into highly curved membranes ^37^. Therefore, for the lipid species studied here, we would expect higher partitioning of PE(18:1|Y), GlcCer(Y), and poly-unsaturated PCs into CCPs, whereas cylindrical lipids such as SM(Y) and saturated PCs would be excluded. In contrast to this hypothesis, we found that the resting curvatures did not correlate with CCP partitioning, suggesting that sorting based on lipid shape is not a plausible mechanism (Figure 4C). This finding is well in line with several studies demonstrating that lipid shape alone is not sufficient for effective lipid sorting (see supporting information for discussion) ^37,38^.

Next, we assessed local phase separation in curved membranes as a possible mechanism ^39–41^. We used the melting temperature *Tm* of the lipid species as a proxy for phase partitioning. According to this hypothesis, highly curved membrane areas should accumulate Ld lipids due to the lower bending rigidity of the Ld phase ^42,43^. This mechanism would result in a negative correlation between lipid melting temperature and CCP partitioning. In contrast, we found a strong positive correlation between melting temperature and partitioning into CCPs (Figure 4C). The long chain saturated PCs, SM, and GlcCer with high *Tm* are enriched while the (poly)unsaturated lipids with low *Tm* are depleted from CCPs. This correlation is surprising as several studies have suggested that Lo lipids with high *Tm* are not trafficked by CME ^10,11,44,45^. Finally, we asked whether the asymmetric distribution of lipids in the plasma membrane impacts partitioning into CCPs. Due to the differences in leaflet surface areas in the curved membrane of the pit and flat surfaces ^33^, we expect that lipids enriched in the exoplasmic leaflet should be partially excluded from the CCP, while lipids enriched in the cytoplasmic leaflet should be partially enriched (Figure 4B). Therefore, we expect a positive correlation between the lipid asymmetry (cytoplasmic/ exoplasmic) and CCP partitioning. Indeed, we found a positive correlation for the asymmetry of all PCs and PE, with the exception of SM (Figure 4C), suggesting that lipid asymmetry is sufficient to explain CCP partitioning for almost all investigated lipids.

Taken together, our correlation analysis suggests that lipid asymmetry of the plasma membrane is the major mechanism underlying lipid sorting in CCPs. It is interesting to note that melting temperature and lipid asymmetry are strongly correlated for glycerophospholipids (Extended Data 10), which could explain the observed positive correlation between melting temperature and CCP partitioning.

**Figure 4:**
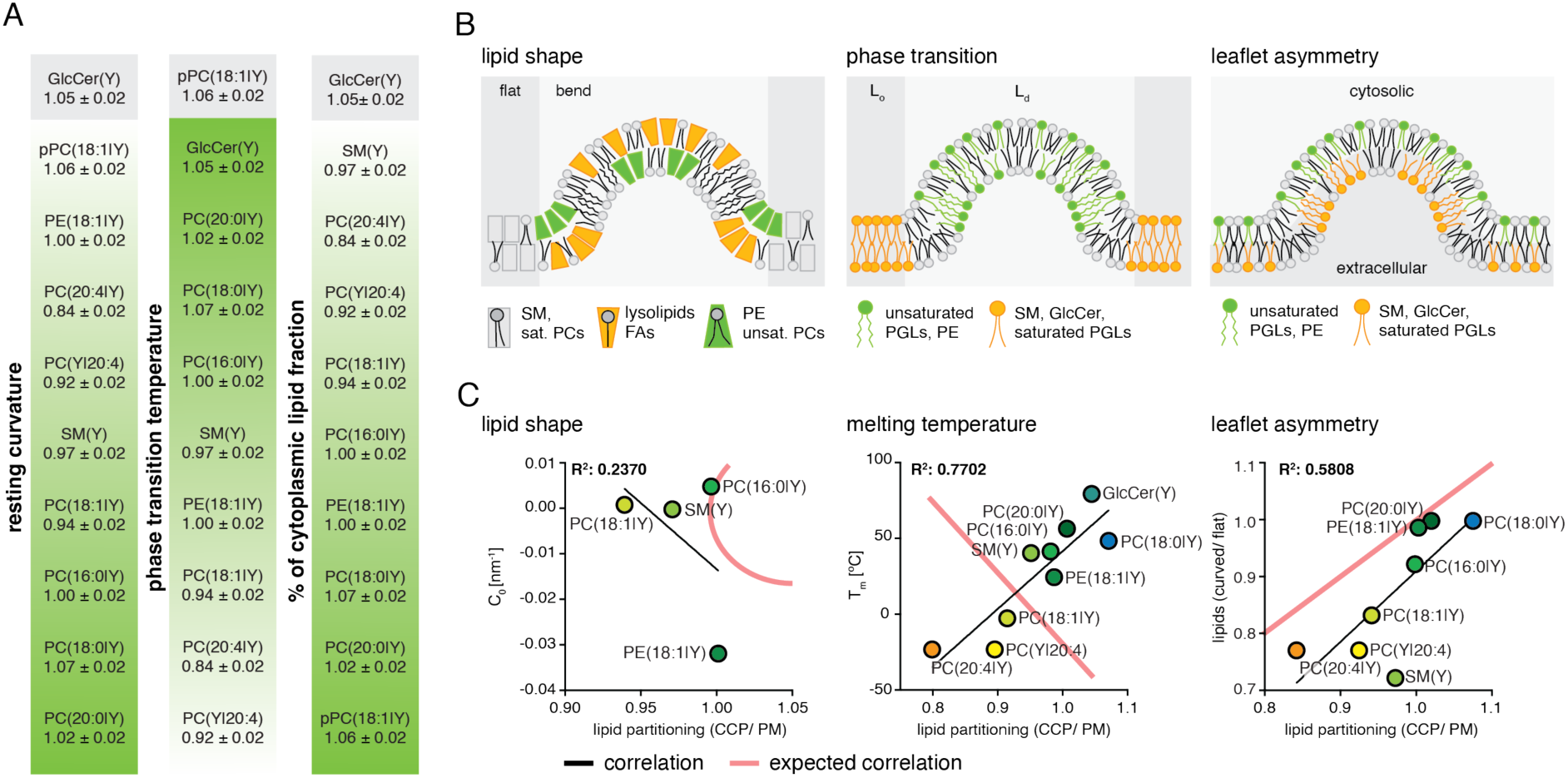
Leaflet asymmetry is sufficient to explain lipid partitioning into clathrin-coated pits. A: Lipids were ordered according to their resting curvature, melting temperature, and transbilayer distribution (cytoplasmic/ exoplasmic) based on reported values from the literature or estimates based on close structural analogs. B: Schemes of possible mechanisms of lipid sorting into clathrin-coated pits are shown. Ld – liquid disordered phase; Lo – liquid-ordered phase. C: The median of absolute lipid densities of late-stage pits is plotted against the respective resting curvature (C0), melting temperature (Tm), and the ratio of estimated lipids in a vesicle over a flat membrane of the same surface area. The values plotted can be found in the supplement 14-16. Only the lipids are plotted for which the corresponding values of C0, Tm, and lipid asymmetry were reported in the literature. The red line is a sketch and indicates the expected trend of the correlation according to the model.

## Discussion

We here introduced super-resolution lipid imaging in combination with mathematical modeling to characterize the extent of lipid sorting during CME. This method allowed us to quantify lipid partitioning into late-stage CCPs. Comparing ten lipid species, we found that lipid partitioning values differed by up to 21% between lipids. A large part of this difference can be explained by the known asymmetric distribution of lipids between plasma membrane leaflets ^33^ in combination with pit geometry. In addition, we found a strong correlation between melting temperature and CCP partitioning. High melting temperature lipids are partitioned more strongly into CCPs, which is in contrast to previous reports ^10,11,44,45^ and the biophysical properties of a curved membrane ^42,43^. However, this observation can be explained at least for the glycerophospholipids by the fact that melting temperature and lipid asymmetry are positively correlated (Extended Data 10). While we cannot exclude other mechanisms, experimentally determined geometric parameters of CCP architecture and lipid-species specific asymmetries are sufficient and plausible to explain the observed glycerophospholipid partitioning effects. Interestingly, sphingomyelin (SM) deviates from this trend, suggesting other mechanisms, such as specific lipid-protein interactions.

These findings suggest that the lipid composition of clathrin-coated vesicles generated during CME is almost exclusively a reflection of the lipid composition of the plasma membrane and its inherent asymmetric lipid transbilayer distribution. Overall, this implies a slight bias for cytoplasmic leaflet lipids to be transported in the retrograde direction. However, this selectivity is small compared to the differences observed during non-vesicular lipid transport ^19,20^. In addition, we previously found that over 80 % of lipid transport occurs via non-vesicular lipid transport mechanisms during retrograde transport. This indicates that lipid transport via CME is a minor portion of the total lipid transport volume ^20^. Therefore, our data suggests that CME does not significantly affect the steady-state composition of the plasma membrane. In contrast, intracellular organelles will shift to a lipid composition more similar to the plasma membrane by CME, which has to be counteracted by other lipid transport processes such as non-vesicular lipid transport.

Taken together, the selective retrograde transport of lipids and proteins appears to be fully decoupled. We propose that the main function of CME is to selectively transport proteins, while the associated lipid transport is largely unspecific. On the other hand, specific lipid transport is mediated by non-vesicular mechanisms. We anticipate that our methodology for quantifying lipid partitioning into cellular membrane compartments of complex geometry will help to understand membrane compartmentalization and the functions of lipid species in cell biology.

## Acknowledgements

AN gratefully acknowledges financial support by the European Research Council (ERC) under the European Union’s Horizon 2020 research and innovation program (grant agreements no GA 758334 ASYMMEM and AURORA). AN and AH acknowledge financial support from the Deutsche Forschungsgemeinschaft (DFG) via the TRR83 consortium. This research was supported by an Allen Distinguished Investigator Award, a Paul G. Allen Frontiers Group advised grant of the Paul G. Allen Family Foundation to AN and AH. LO gratefully acknowledges financial support from the Czech Science Foundation (Project No. 24-10736S). We thank the following facilities at MPI-CBG Dresden for the support: Light Microscopy Facility. We thank Lisa Redlingshöfer and Ori Avinoam for insightful discussions and valuable advice. We are grateful to the Drubin lab for the kind gift of the SK-MEL2 dymanin2 GFP reporter cell line.

## Author contributions

KB and LO synthesized lipid probes. HML developed the STED lipid imaging workflow. HML and SCN acquired the STED dataset. HML analyzed fluorescence imaging data. LM, WYY, and CM designed and built mathematical models. HML, SCN, SK, and JMIA prepared samples. HML, AH, and AN designed the project. AH, CM, and AN supervised research. HML, AH, and AN wrote the manuscript. All authors read and commented on the manuscript.

## Data Availability

Data deposition: All software codes used for analysis and data have been deposited on the Max Planck Institute of Molecular Cell Biology and Genetics repository and are available for the reader at https://doi.org/10.17617/3.GLIVDU and upon request from the corresponding authors.

## Extended Data

**Extended Data 1.**
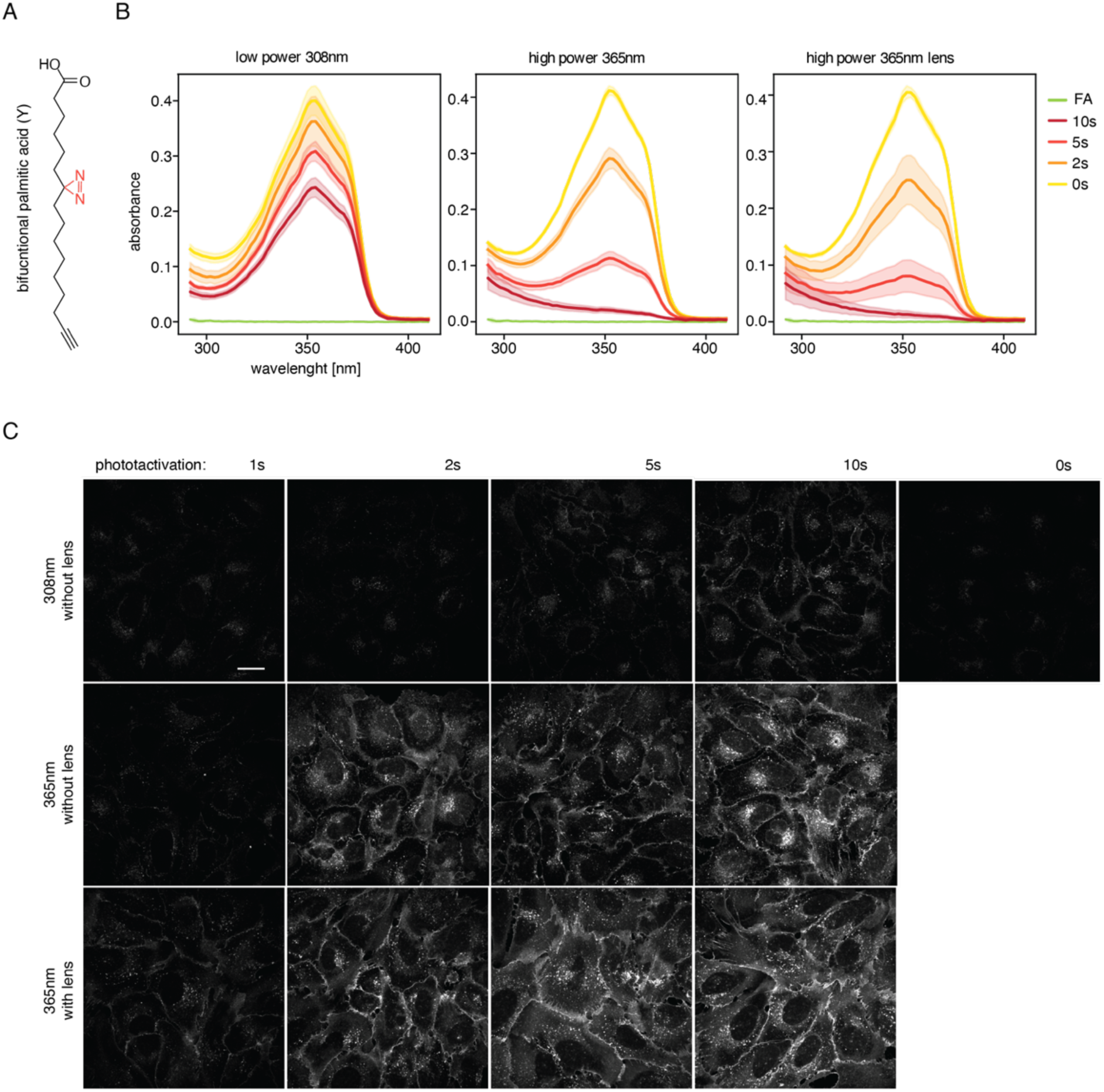
Characterization of the diazirine photo-crosslinking efficiency. A: The chemical structure of the previously reported ^20^ bifunctional palmitic fatty acid (Y) is shown. B: The diazirine spectrum of Y is measured at 100mM concentration in DMSO. Spectra were measured after different exposure times of 0s (yellow), 2s (orange), 5s (red), and 10s (deep red) with a recently reported 308nm LED ^20^ compared to 2 high power LEDs at 365nm with and without a collimating lens. The spectra of the native fatty acid (FA, green) lack a diazirine peak. Photoactivation experiments were performed in triplicates. Mean values and standard deviation are depicted. C: U2OS wildtype cells were loaded for 4min with bifunctional SM, UV activated for different time intervals and with different LEDs. Signal intensities are scaled equally for all images. Scale bar is in 20µm.

**Extended Data 2.**
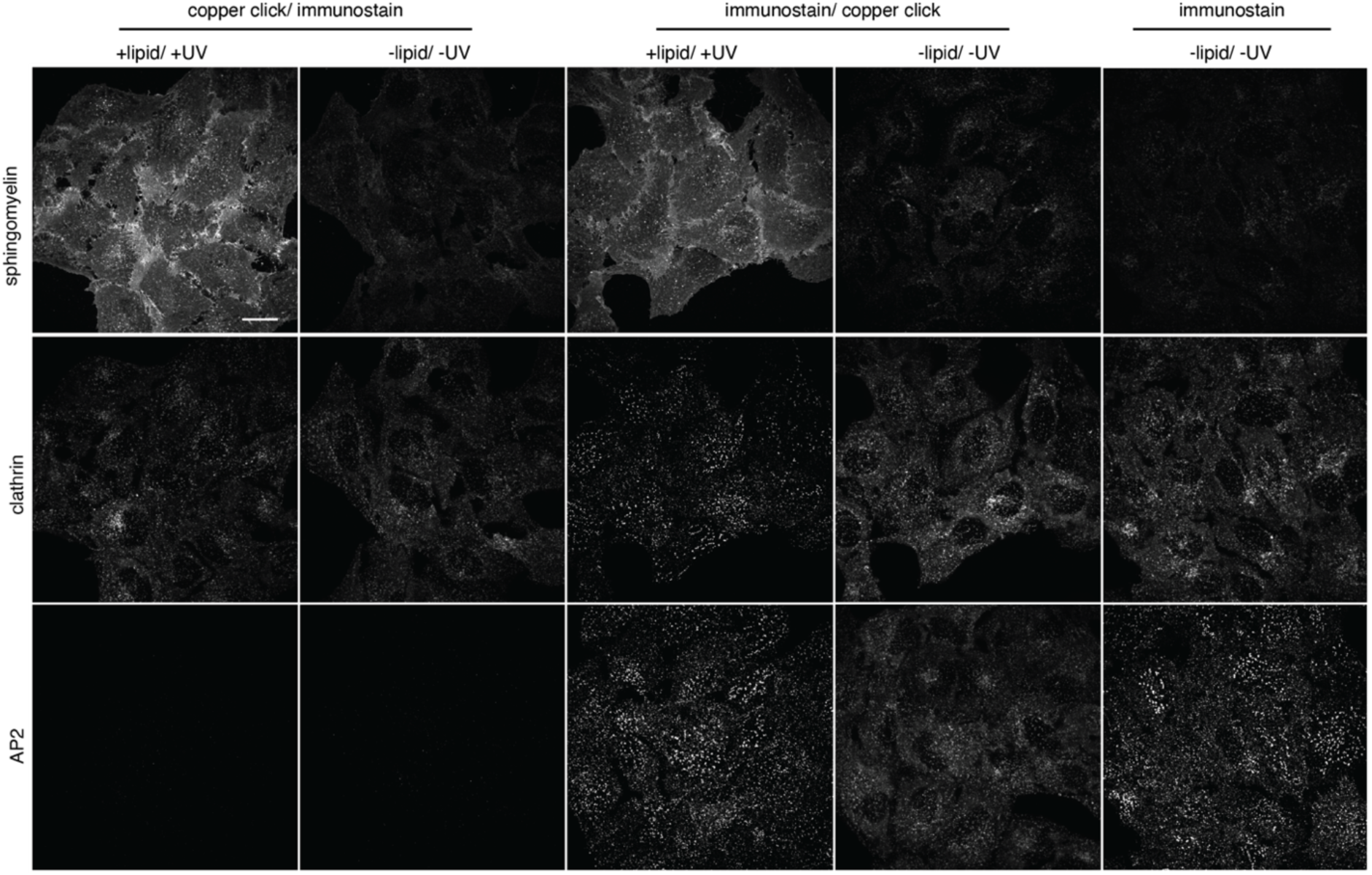
Copper-catalyzed click chemistry can prevent immunolabelling. U2OS wildtype cells were fed for 4min with bifunctional sphingomyelin, photoactivated, chemically fixed, and permeabilized. Samples were then either first stained by copper-catalyzed click followed by immunolabelling (copper click/ immunostain) of clathrin and AP2 or first immunolabelled and then stained by copper-catalyzed click (immunostain/ copper click). As a control for the immunolabelling efficiency, a sample was only immune-labeled but not stained by copper-catalyzed click (immunostain).

**Extended Data 3.**
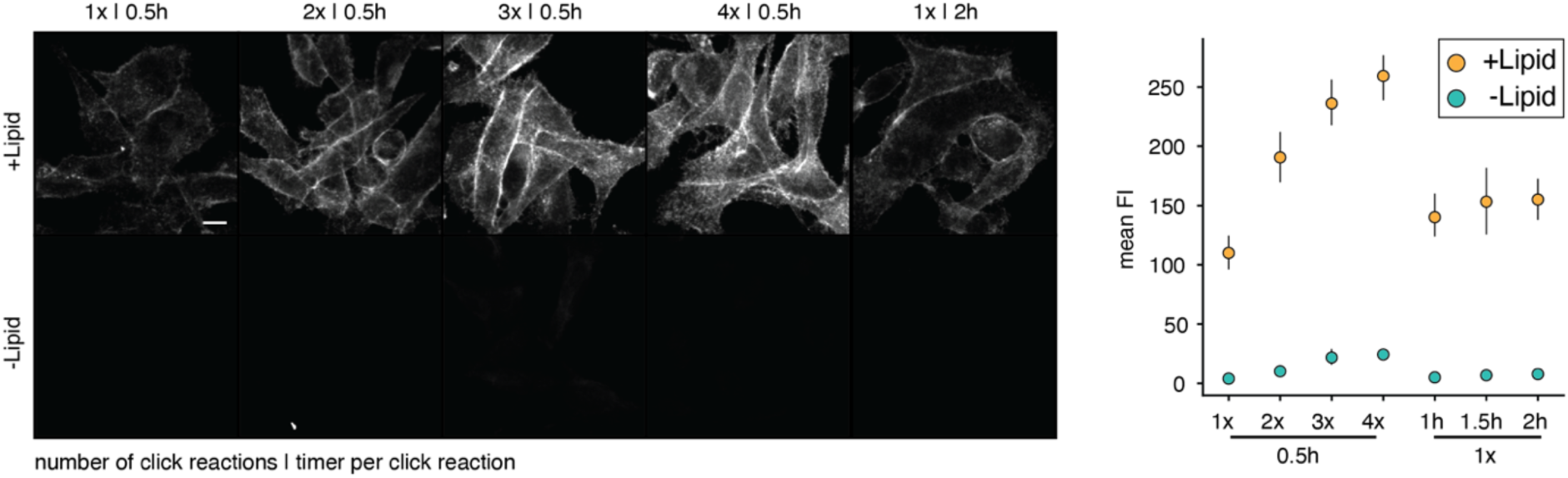
Multiple copper-catalyzed click reactions increase lipid signal. SKMEL wildtype cells were loaded with PC(16:0/Y) for 4min. Samples were stained by CuAAC 1, 2, 3, or 4 times for a duration of 30min per stain or 1 time for the respective prolonged duration of 1h, 1.5h, or 2h. Scale bar: 10µm. Fluorescent images were background corrected and mean intensity values were determined. Error bars represent the standard deviation. The experiment was performed in triplicates.

**Extended Data 4.**
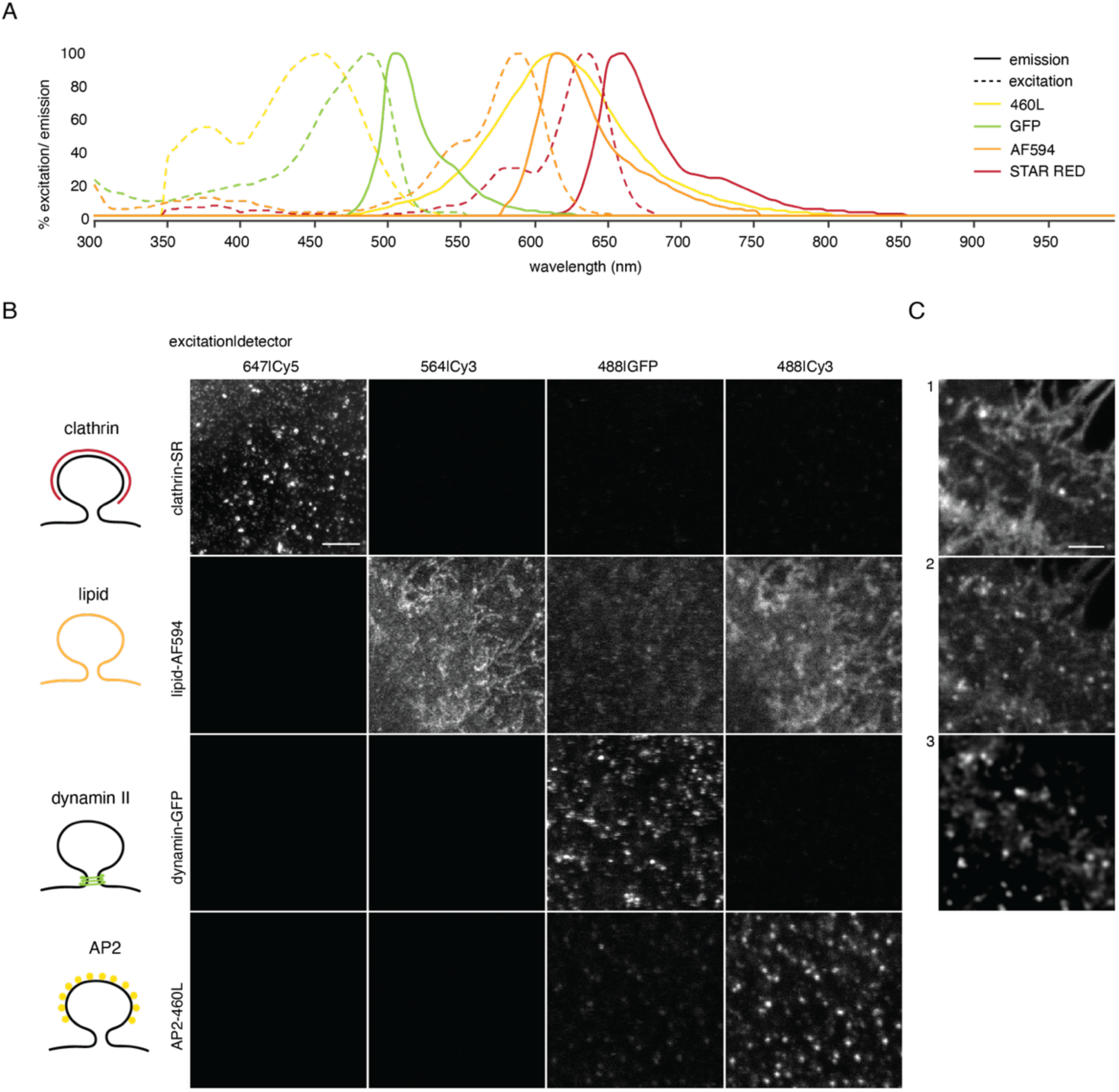
Bleed-through correction by linear de-mixing. A: Spectra of the used fluorophores abberior 460L, GFP, AlexaFluor594, and abberior STAR RED generated with the FluoroFinder Spectral Viewer. B: SK-MEL cells with a single label, either clathrin – STAR RED (row 1), lipid – AlexaFluor594 (row 2), dynamin-2-GFP (row 3), or AP2 – 460L were imaged with all four imaging channels to determine bleed-through. Excitation lasers used were 647nm (clathrin), 564nm (lipid), and 488nm (dynamin-2, AP2). Emission was collected in the Cy5 (clathrin), Cy3 (lipid. AP2), or GFP (dynamin-2) channel. C: The confocal image of the AF594 (1) and the confocal image of the 460L (2) are shown. After linear de-mixing (3) the background from the 460L images was removed. All scale bars are shown in 2µm.

**Extended Data 5.**
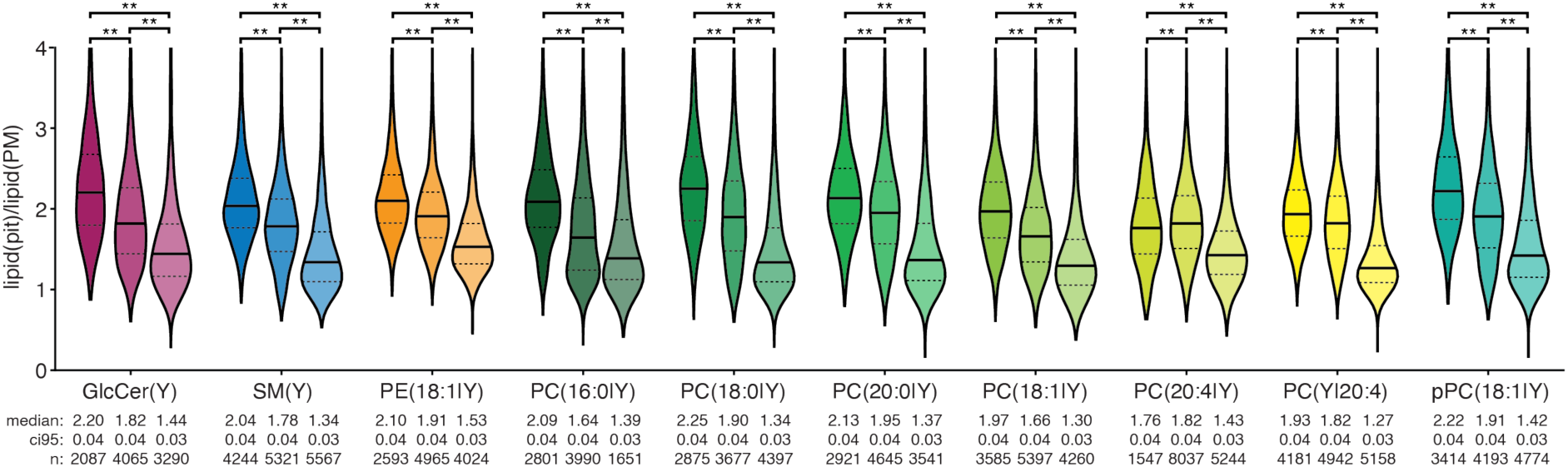
Lipid enrichments of clathrin-only, early-stage, and late-stage CCPs. Violin plots of the distributions of lipid pit enrichments over all measured individual pits are plotted for the different lipids and structure classes. The median (solid line) and quartiles (dotted line) are shown. For each lipid the structure class distributions differ significantly as assessed by a two-sided randomized permutation analysis (** indicates p-values below 0.001). The experiment was performed for 3 biological replicates.

**Extended Data 6.**
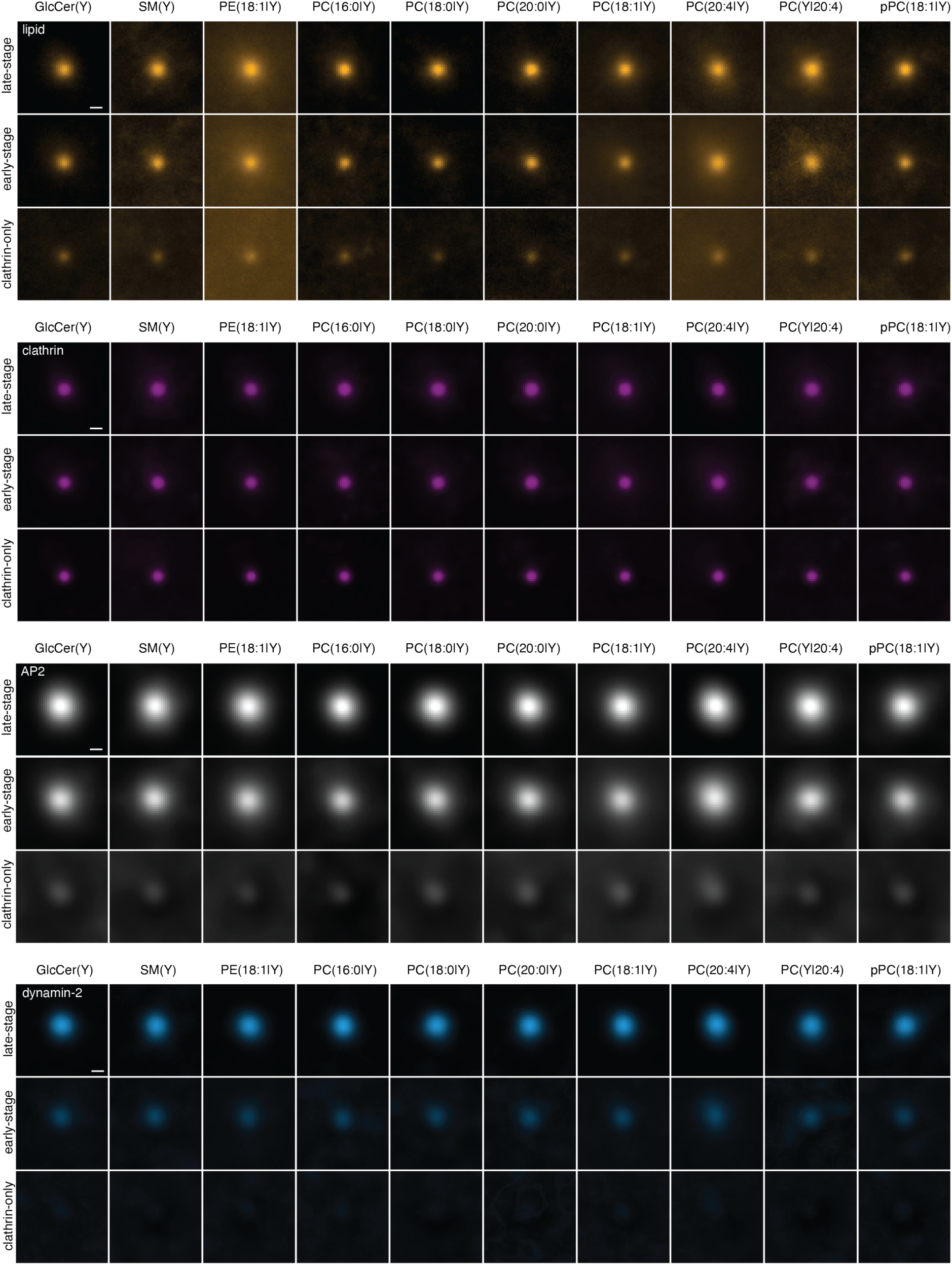
Averaged images of CCPs of all lipids and CCP stages. All fluorescent images for all lipids species of CCPs (square of 1.5µm x 1.5µm) were centered to the centroid of the clathrin mask and averaged for clathrin only, early- and late-stage pits. The lipid (orange), clathrin (purple), AP2 (grey), and dynamin-2 (blue) channels are displayed separately. Scale bar: 100nm. Within one channel all images are scaled to the same brightness and contrast. The average images for early- and late-stages for PC(16:0|Y) are also displayed in Figure 2C and all late-stage lipid images are also displayed in the main Figure 3B.

**Extended Data 7.**
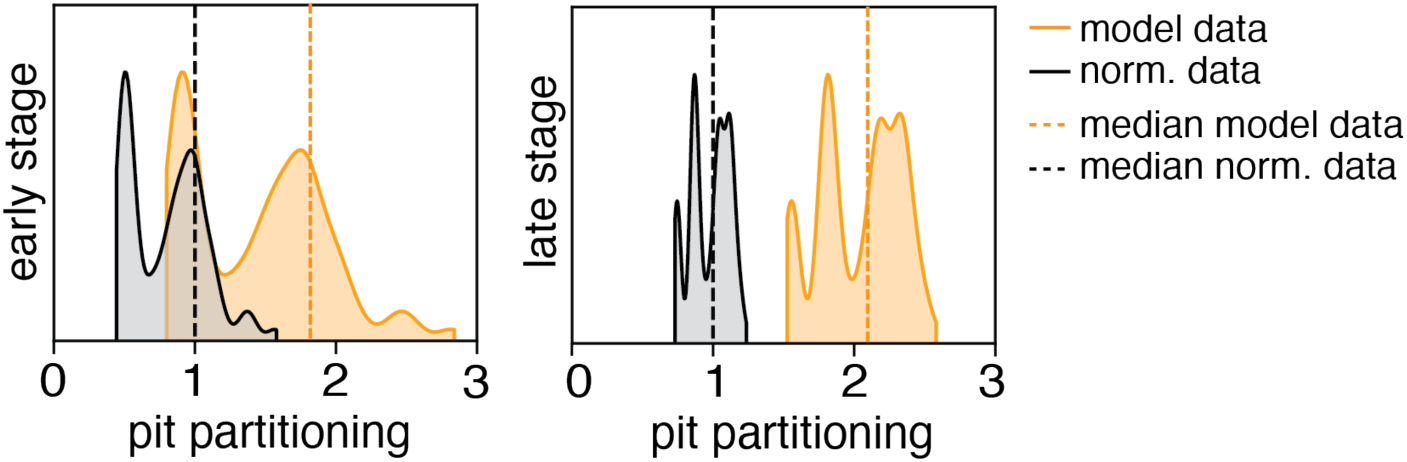
Comparison of the model data with the geometry model normalized to the median of the model. The pit partitioning distributions of the model (orange) of early and late pits are overlayed with the respective distribution normalized to the median of the model (grey). The black and orange dotted lines indicate the median of the model before and after normalization respectively.

**Extended Data 8.**
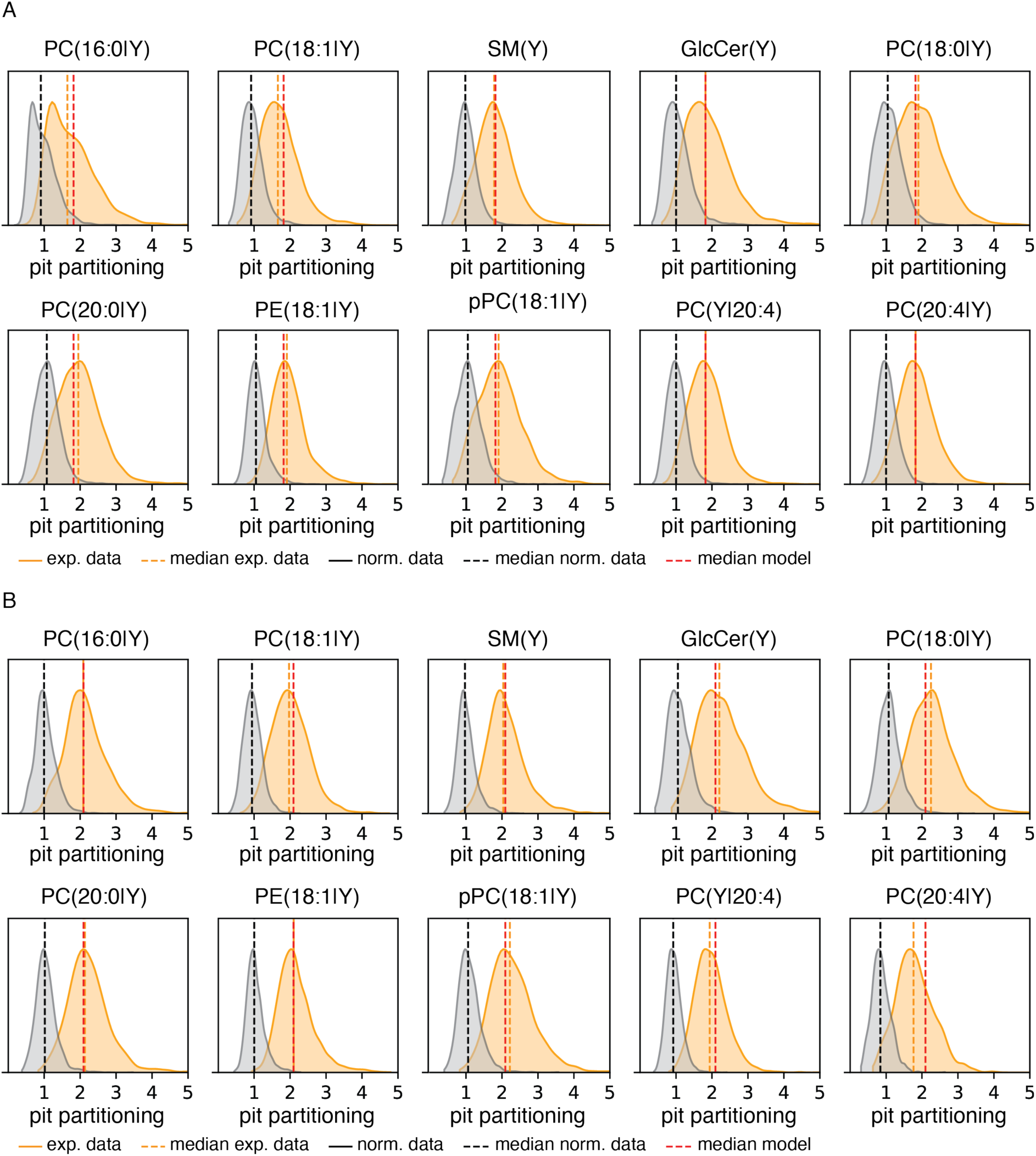
Comparison of the non-normalized and normalized experimental data. The experimentally determined pit partitioning distributions (orange) of early (A) and late pits (B) are overlayed with the respective distribution normalized to the median of the model (grey). The lipid species is indicated on top of each plot. The black, red, and orange dotted lines indicate the median of the model, experimental data, and normalized experimental data respectively.

**Extended Data 9.**
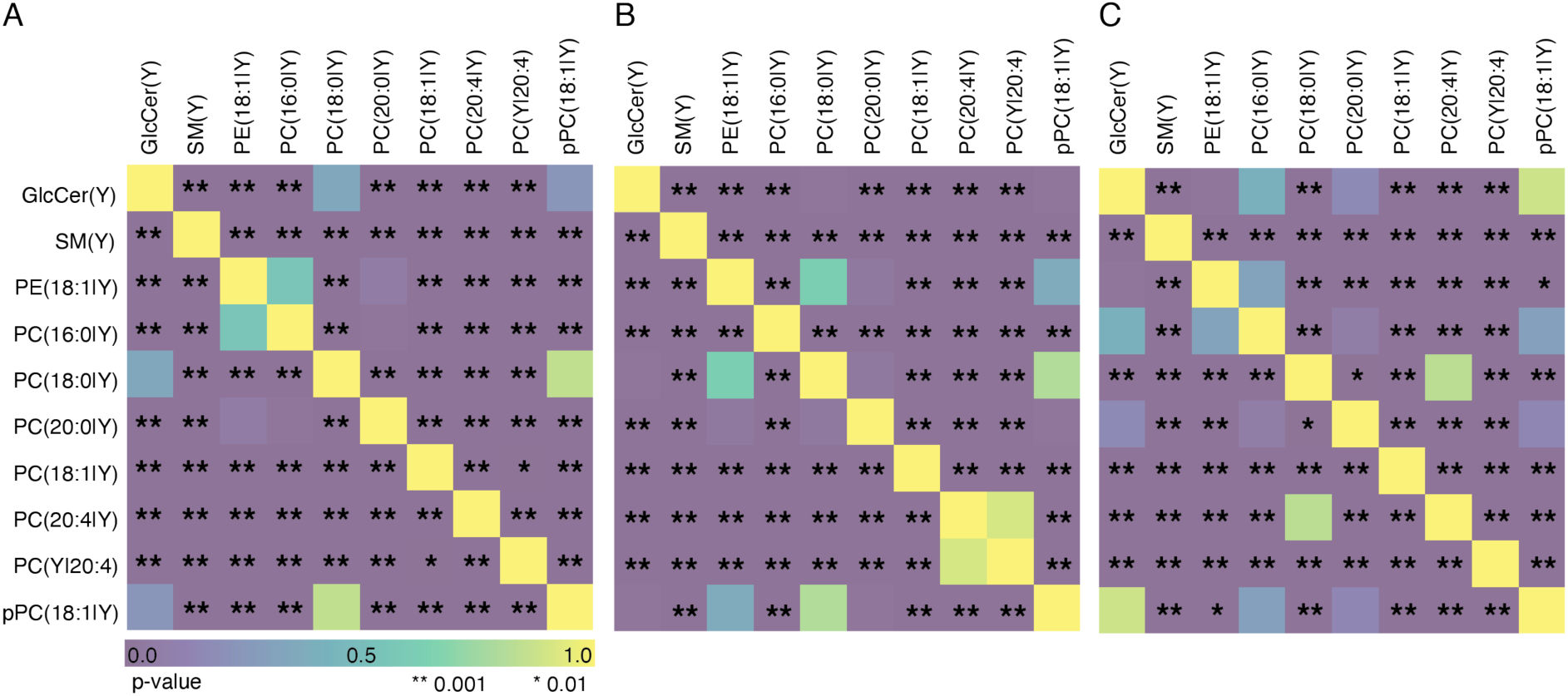
Two-sided randomized permutation test for the lipid enrichment distributions. Significance between lipid pit enrichments of late (A) and early pits (B) and clathrin-only (C) structures. P-values per lipid pair were determined by a two-sided randomized permutation analysis ( https://rasbt.github.io/mlxtend/user_guide/evaluate/permutation_test/). P-values are color-coded with * marking values below 0.01 and ** marking values below 0.001. The significance between lipid pit enrichments of late-stages (A) is also shown in the main figure 3D.

**Extended Data 10.**
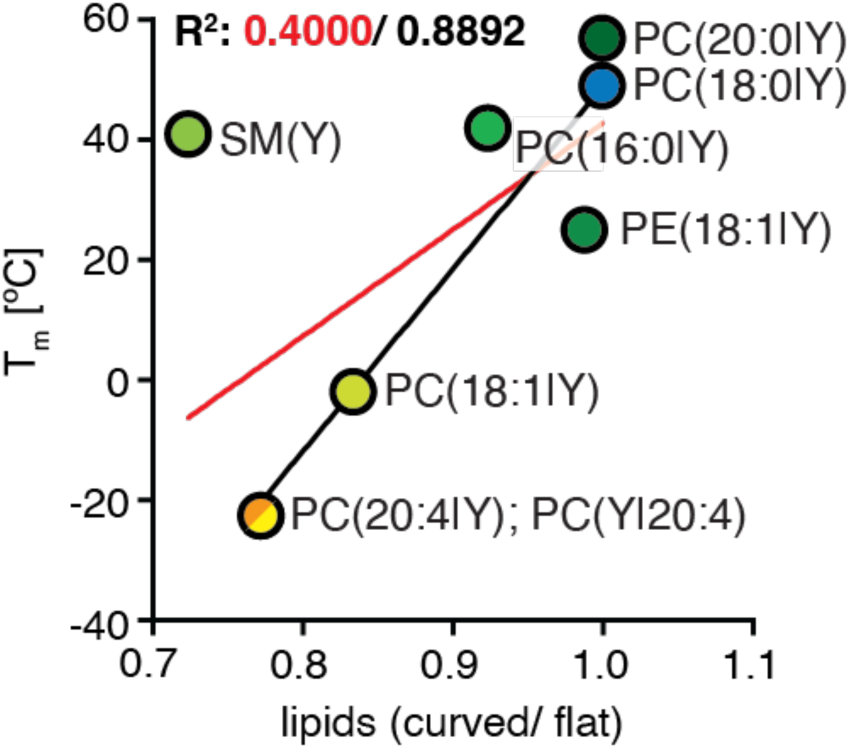
Correlation between lipid asymmetry and melting temperatures. The melting temperatures (Tm) of different lipid species are plotted against the respective ratio of estimated lipids in a vesicle over a flat membrane of the same surface area. The values plotted can be found in the supplement 14 and 15. Only the lipids are plotted for which the corresponding values of Tm and lipid asymmetry were reported in the literature. The red line indicates the linear regression fit between all lipids, and the black line indicates the same fit but excluding SM(Y). The corresponding R^2^ values are indicated on top in red and black, respectively.

