## Supplementary Information for "Quantification of lipid sorting during clathrin-mediated endocytosis"

**Supporting Information for**
**Quantification of lipid sorting during clathrin-mediated endocytosis "**

H. Mathilda Lennartz, Susana da Costa Nunes, Kristin Böhlig, Sascha M. Kuhn, Lior Moneta, Wan Yee Yau, Lukáš Opálka, Juan M. Iglesias-Artola, Carl D. Modes, Alf Honigmann and André Nadler

**Data Availability**
Codes and data can be found in the following repository: doi:10.17617/3.GLIVDU.

**This PDF includes**

- Materials and Methods
- Supplementary Data 1 to 4
- Supplementary Figures 1 and 2

### Methods and Materials

#### LED characterization

LEDs were custom built in-house. The following LEDs were tested for their power output and crosslinking efficiency: a 308nm LED without lens (PKF-50, UVB LED 308nm +/ -5nm # 3049140, Laser Components) as previously reported [1] and a high power 365nm LED with (UVB LED 365nm +/ -10nm, #VC2X2C45L9-365, violumas) and without lens (UVB LED 365nm +/ -10nm, #VC2X2C45LX-365, violumas). The power was measured using a PM400 Optical Power Meter from Thor Labs with a S175C Sensor from Thor Labs. The power stated is the maximum power measured over a 10s exposure time. The power was measured for the wavelength specific for each LED (either 308nm or 365nm). Crosslinking efficiency was determined by 10s, 5s, 2s, and 1s irradiation of a 100mM bifunctional palmitic fatty acid in DMSO solution. UV irradiation was performed in 96-well plates with glass bottoms (greiner, #655891). Spectra of the bifunctional palmitic fatty acid were measured with a TECAN SPARK 20M in 96-well black plate with a plastic bottom (greiner, #655090) and using no lid. Crosslinking efficiency in U2OS wildtype cells was determined by 10s, 5s, 2s, and 1s irradiation of bifunctional sphingomyelin with the respective LED.

#### Cell culture

SK-MEL2 dynamin-2-GFP and SK-MEL2 wildtype cells (kind gift from D. Drubin, described in Doyon et al. [2]) were cultured in DMEM/ F-12 + GlutaMAX (gibco, #10565-018) medium supplemented with 20% fetal bovine serum and 100U/ml Penicillin-Steptomycin at 37°C with 5% $CO_2$ in 75nm$^2$ flasks. Cells were split every 3-4 days using 0.05% Trypsin, ensuring they never reach a confluency of 100%. Cells were used up to passage 15. For experiments cells were seeded 24h before the treatment into fibronectin bovine plasma (sigma, #F4759-2MG) coated 96-well plates with glass bottom (greiner, #655891) at a density of 20,000 cells per well. Fibronectin coating was done at 37°C for 30min before cell seeding at 10µg/ml in phosphate-buffered saline (PBS).

#### Liposome preparation and lipid loading

A suspension of 1.5mM bifunctional lipid (azide or alkyne), 0.75mM 1-palmitoyl-2-oleolyl-glycerol-3-phosphocholine (Avanti, #26853-31-6) and 0.75mM cholesterol (sigma, CAS: 57-88-5, #C8667-5G) in PBS was prepared. Liposomes were made from this suspension using an Avanti Mini-Extruder with 0.1µm polycarbonate membranes (Avanti, #610005-1Ea), extruding a minimum of 21 times. Liposomes were stored for up to 2 weeks at 4°C. Liposomes and alpha-methyl-cyclodextrin (Bio-reagent, #CDexA-076/BR, CAS: 699020-02-5) were dissolved in serum-free medium to a final concentration of 0.5mM and 4mM, respectively. This mix was incubated for at least 30min at 37°C before loading on cells. Before lipid loading cells were gently washed 3 times with serum-free medium. Lipids were loaded for 4min at 37°C to cells for STED imaging and 30s for imaging by Spinning Disc microscopy. Cells were washed 3 times in serum-free medium, followed by immediate photoactivation (unless stated otherwise) using a custom-built high-power 365nm LED (UVB LED 365nm +/- 10nm, #VC2X2C45L9-365, violumas).

**Cell fixation and Immunostainings**

Samples were fixed immediately after photoactivation for 20min at room temperature using a fixation buffer of 3% PFA, 10mM MES, 150mM NaCl, 5mM EGTA, 5mM d glucose, 5mM $MgCl_2$, pH 7.2. Fixed cells were washed 3 times with 100mM glycine in PBS and permeabilized using 0.1% Triton in PBS for 30min at room temperature. Fixed and permeabilized samples were blocked at room temperature for 1h in blocking buffer composed of PBS supplemented with 2% Bovine serum albumin (BSA, Sigma, #A7030-10G). Primary Antibodies were incubated for either 1h at room temperature or overnight at 4°C in blocking buffer. Samples were washed 3 times for 15min in blocking buffer at room temperature and incubated with the respective secondary antibody for 1h at room temperature in blocking buffer. Samples were washed 3 times for 15min in blocking buffer at room temperature and kept in PBS. Samples were always immunolabelled before staining by click chemistry. Table 1 shows all primary and secondary antibodies used.

Table 1| Antibodies

| Type | Target protein | animal | provider | Product nµber | Buffer system | dilution |
|---|---|---|---|---|---|---|
| prim | Clathrin light chain | rabbit | proteintech | 10852-1-AP | 2% BSA in PBS | 1:200 |
| prim | AP2 [AP6] | mouse | Invitrogen | MA1-064 | 2% BSA in PBS | 1:200 |
| prim | EEA1 | mouse | BD Bioscience | 610457 | 2% BSA in PBS | 1:200 |
| sec | STAR 460L anti mouse | goat | Abberior | ST460L-1001 | 2% BSA in PBS | 1:100 |
| sec | STAR RED anti rabbit | goat | Abberior | STRED-1002 | 2% BSA in PBS | 1:100 |
| sec | AF488 anti mouse | goat | Invitrogen | A11001 | 2% BSA in PBS | 1:1000 |

**Copper-catalyzed click reaction**

Samples were washed and equilibrated in 100mM HEPES and stained using copper-catalyzed click reaction by incubation at 37°C with a click mix of 2µM AF594 picolyl-azide dye (jena bioscience, CLK-1296-1), 0.1mM CuSO4, 5mM ascorbic azide and 0.5mM THPTA (jena bioscience, CLK-1010-1G) in 100mM HEPES at pH 7.3. Samples were stained 3 times with fresh click mix for 30min each.

**Spinning disk imaging**

Images were acquired on an Olympus IX83 microscope, equipped with a Yokogawa CSU-W1 SoRa unit, an ORCA-Fusion from Hamamatsu, and an ORCA-Flash 4.0 V3 digital CMOS camera. A 100x immersion oil objective (Olympus UApoN OTIRF) was used for imaging. A 488nm laser line was used to image EEA1 at 10% power and a 561nm laser line was used for the lipid at 50% laser power with their respective filter sets. All images for the lipid transport dataset were acquired using identical settings. Cells were imaged using 17-slice z-stacks, with 0.5µm distance in z between frames. To ensure that all images were acquired at the same starting plane, the Olympus TruFocus Z-drift compensation system was used. Each frame was exposed for 100ms. The pixel size was 65nm.

**STED microscopy**

Images were acquired using a commercial confocal infinity-line STED microscope (Abberior Instruments, Göttingen, Germany) equipped with a pulsed laser excitation (640nm, 560nm, 460nm, 40MHz), beam scanning module (line frequency 3kHz), and single photon counting APD detectors. As the objective, a 100x/1.49 NA oil (Olympus) was used. A 775 nm, 40MHz pulsed laser (Katana HP, 3W, 1ns pulse duration, NKT Photonics) was applied for the depletion of the 560nm and 640nm channel. A pinhole size of 60µm was used. Images were acquired at a pixel size of 30nm. The star red dye, AF594, star L460, and GFP were excited at 640 nm, 561nm, 488nm, and 488nm and detected in a range of 685nm | 70nm, 605nm | 50nm, 525nm | 25nm respectively. The system was operated using the Imspector software (v.16.2.8415).

**Image Analysis**

Image and data post-processing was performed using python [3] , Ilastik [4], and ImageJ/ Fiji [5,6]. The python version 3.8, ImageJ/ Fiji version 2.14.0/1.54f, and Ilastik version 1.4.0-OSX were used.

**Linear demixing**

The signal from the AlexaFluor594 fluorophore showed bleedthrough into the LS channel and was corrected by linear demixing. The correction factor was determined from negative control samples only stained with AlexaFluor594 by division of the long stokes shift convocal signal (bleedthrough) over the 594 convocal signal using Fiji [6]. Following, both the confocal image of the 594 channel and the confocal long stokes shift channel were blurred by applying a gaussian filter. The product of the correction factor and the blurred 594 images were subtracted from the respective blurred long stokes shift images using the self-written script *240724_linear_demixing.py* .

**STED image segmentation analysis**

The linear demixed confocal AP2 images and confocal dynamin-2 images are segmented using conventional thresholding (threshold yen, skimage). The STED clathrin images were segmented by upsizing images 5fold (cubic interpolation, resize, cv2), rescaling intensity values from 0 to 1, and segmenting (threshold otsu, clesperanto) and labeling (voronoi otsu labeling, clesperanto) using napari. Label images were downsized to the original dimensions (nearest interpolation, resize, cv2). Next, the raw STED lipid images were background corrected. Using the clathrin labels and the background corrected lipid images, the mean fluorescence lipid signals, area, and centroid coordinates were determined per pit (regionprops.measure, skimage). Segmented clathrin pits were assigned as unidentified structures if they did not overlap with both the AP2 and dynamin-2 masks. Pits overlapping with only AP2 were classified as early-stage pits, and pits overlapping with both masks as late-stage pits. Pits below a size of 5 pixels were considered as noise and filtered out. Next, using the centroid coordinates of each pit, a square of 1.5µm x 1.5µm (referred to as ROI) was defined around this center. The mean lipid fluorescent signal of each ROI was determined from background-corrected lipid images where the areas without cells, high-intensity filopodia, and regions of clathrin masks were set to *nan* to be excluded from the mean intensity of each ROI. The pit lipid enrichment was calculated by division of the fluorescent mean pit/ mean ROI. For each lipid type and each pit class, the mean pit lipid enrichment was calculated with the respective standard deviations (std) and total number of analyzed pits. The significance values between different lipid mean pit lipid enrichment per pit class were calculated using a two-sided randomized test (https://rasbt.github.io/mlxtend/user_guide/evaluate/permutation_test/). Clathrin pit area densities were determined for each pit class per image by taking the ratio of the area (in pixel number) for all pits over the total cell area (in pixel number). Clathrin pit number densities were determined for each pit class per image by taking the ratio of the number of all pits normalized over the total cell area (in pixel number). Data was plotted using seaborn. The code can be found in the *240724_CCP_segmentation_analysis.py* file.

**Ilastik models**

We used the 2-stage Autocontext pixel classification workflow of Ilastik [4] for segmenting early endosomes co-stained with EEA1 in the 488 channel. The training was performed in 2D and using different frames of the Z-stack.

**EEA1 segmentation and analysis**

Image z-stacks were analyzed by single images. The EEA1 label was segmented using the Ilastik script *EEA1_SKMELwt.ilp* . Lipid images were background corrected using the mean fluorescent value of the no lipid control. The total fluorescent lipid intensity counts were determined per image, not counting cell-free areas. The fluorescent lipid intensity counts were determined for the pixels overlapping with the EEA1 mask (EEA1 intensity counts). Per image z-stack, the sum of the total lipid intensity counts and the EEA1 intensity counts was taken. The ratio of the EEA1 intensity counts over the total lipid intensity counts per image z-stack was considered the total lipid amount in early endosomes in %. This analysis was performed using the self-written python script *270724_EEA1_SKMEL_segmentation.py* .

**Clathrin-coated pit modeling**

Pit geometry modeling was performed by sampling points on a simplified pit geometry. We assumed uniform point distribution on the pit and the membrane surrounding it with point density $\rho$. We used a cut sphere to represent the pit and a catenoid to represent the neck. The radius and opening angle of the cut sphere along with the height and width of the catenoid, defined the pit geometry. Electron microscopy measurements of CCPs from Avinoam et al. [7] were used to generate biologically accurate pit shapes. We sampled $n_{pit} \sim \mathcal{N}(N_{pit}, 0.1N_{pit})$ points on the cut sphere and the catenoid, where $N_{pit} = A_{pit} \times \rho$ and $A_{pit}$ is the pit's surface area. Points were allocated between the cut sphere and the catenoid based on their relative surface areas. Standard uniform sampling on a sphere was utilized on the cut sphere, while a geometrical sampling algorithm developed by Modes et al. [8] was used for the catenoid. We simulated 1000 early-stage pits and 1000 late-stage pits using the pit angle distribution from Mund et al. [9] to achieve the proper shape distribution of the pits. Due to observed inconsistencies between the pit and neck parameters in a small subset of EM data, we needed to exclude certain generated pits from our analysis. We use a threshold $\delta = 1[nm]$ on the difference between the spherical cap opening radius $r_s$ and the catenoid opening at the reported height $r_c$ as defined in equation 1. Any pit with $\Delta r > \delta$ is excluded from the analysis. Our threshold excludes (percentage of pits by the chosen delta)of the pits.

$$\Delta r = |r_s - r_c| \quad (1)$$

**STED image simulation and segmentation**

Using the coordinates of the simulated point emitters from the clathrin-coated pit modeling, STED images were simulated with a similar pixel size of 30, 30, 500nm (x, y, z) as the experimental imaging data. The size of the STED microscope point spread function (PSF) was determined by imaging 20 nm Crimson beads and measuring the full width at half maximum (FWHM) in the XYZ axis. Simulated point emitters were convolved with a gaussian PSF with FWHM of 80, 80, 500nm (x, y, z) according to the experimentally determined PSF parameters using custom Python code to generate STED images of the 3D models. The generated images were cropped to a 50 pixel x 50 pixel array to remove an edge effect of lower intensities at the image edges caused by the convolution. For the segmentation of the simulated images, a line intensity profile of the experimental data was generated for all early and late-stage pits. The profile was taken through the centroid of the clathrin mask, reading out both the lipid signal intensity and mask intensity. From the individual profiles, we determined the average width of the mask for early and late-stage pits, as well as the respective average intensity profile of the lipid signal using the python script *CCP_average_lineprofile_all_pits.py* . Average intensity profiles were normalized from 0 to 1 and the normalized intensity cutoff for segmentation in % was determined using the average segmentation mask width. The resulting cutoff was used on the intensity profiles of the individual simulated images to determine individual widths of segmentation masks for the model. Masks were generated as symmetric circles. The ratio between the mean fluorescent intensity within the mask and the mean intensity outside the mask was defined as the apparent partitioning factor. The image simulation and analysis of simulated images was done using the custom python codes *Simulate_Pit_STEDImages_early.py* and *Simulate_Pit_STEDImages_late.py* .

**Correlations and Exponential fits**

The linear correlations and one-phase association exponential fits were performed using GraphPad Prism version 10.4.0 for macOS, GraphPad Software, Boston, Massachusetts USA, www.graphpad.com. For the linear correlation analysis values of melting temperatures, resting curvature, and lipid asymmetry were determined from the literature (see Supplement 1 -3), and these values were correlated against the experimentally determined pit partitioning values. For this analysis, we assume that PC(20:4|Y) and PC(Y|20:4) have the same values for Tm, resting curvature, and lipid asymmetry. For the lipid asymmetry, we assume that PC(20:0|Y) is localized fully to the cytosolic side based on the reported values of PC(18:0|Y). We estimated the melting temperature of PC(20:0|Y) based on the values of PC(16:0|Y) and PC(18:0|Y). For pPC(18:1|Y) and GlcCer(Y) no literature values were found, and thus, these lipids were excluded from the correlation analysis. For the one-phase association, exponential fits of the values for the mean lipid amount in early endomes were determined for all lipids and all time points as discussed in the EEA1 segmentation and analysis. We assumed that at timepoint 0 min, no bifunctional lipid content is localized in the early endosomes. Following a non-linear regression (curve fit) for one-phase association was performed to determine the rate constants. To determine the values of amplitudes, we performed the same fit with the constraint that the rate constant should be the same for all traces.

#### Synthesis and Characterization of new Chemical Probes

All chemicals were obtained from commercial sources (Acros, Sigma-Aldrich, TCI chemicals, Avanti Polar Lipids, Alfa Aesar, Roth, Fluka or Merck) and were used without further purification. Solvents for flash chromatography were obtained from VWR and Penta Chemicals and dry solvents from Sigma. Deuterated solvents were obtained from Deutero GmbH, Karlsruhe, Germany. TLC was performed on precoated plates of silica-gel (Merck, 60 F254) using UV-light (254 nm or 365 nm) or staining solution of phosphomolybdic acid in EtOH (3 g phosphomolybdic acid in 100 ml EtOH) for analysis. Flash column chromatography was performed using silica gel from Merck (silica 60, grain size 0.040-0.063 mm) with a pressure of 1 bar. $^{1}$H-, $^{13}$C- and $^{31}$P-NMR spectra were measured on 400 MHz Advance$^{TM}$ III HD Nanobay Bruker spectrometer. Chemical shift of $^{1}$H- and $^{13}$C-NMR spectra are referenced indirectly to tetramethylsilane. J values are given in Hz and chemical shifts in ppm. Splitting patterns are mentioned as follows: s, singlet; d, doublet; t, triplet; q, quartet; m, multiplet; $m_c$, centered multiplet. $^{13}$C-NMR spectra were broadband hydrogen decoupled. Mass sepctra (ESI) were acquired using a QExactive instrument (Thermo Fisher Scientific) equipped 660 with a robotic nanoflow ion source.

#### Synthesis of bifunctional lipid probes

SM(Y), PE(18:1|Y), PC(16:0|Y), PC(18:1|Y), PC(20:4|Y), PC(Y|20:4) and the C16 bifunctional fatty acid were synthesized according to recently published protocols [19,20]. The analytical data was in accordance with those reported. Synthetic procedures and analytical data for new compounds are reported below.

#### Synthesis of PC(18:0|$Y_{16}$)

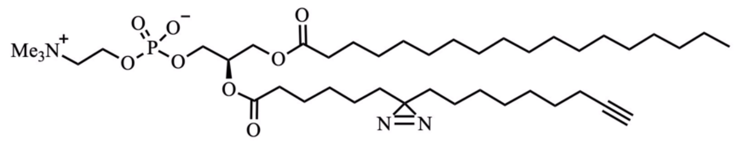

A solution of 31.0 mg of the bifunctional C-16 fatty acid ($Y_{16}$) (111 µmol, 1.2 eq.) in 1 ml $CDCl_3$ was treated with 40.0 mg EDC·HCl (209 µmol, 2.2 eq.) and 5.5 mg DMAP (45 µmol, 0.50 eq.). After 10 min, the solution of the activated fatty acid was added to a solution of 48.6 mg 1-*O*-steaoryl-2-lyso-PC (92.8 µmol, 1.0 eq.) in 2 ml dry DMF and 1 ml $CDCl_3$. The reaction mixture was stirred overnight at room temperature. After removing the solvents under reduced pressure, the product was purified by flash chromatography ($CHCl_3$/MeOH/$H_2O$ 70:30:2). PC(18:0/$Y_{16}$) was isolated as a yellowish oil.

$^1$H NMR (400 MHz, $CD_3OD$) δ = 5.24 ($m_c$, 1H), 4.44 (dd, $J$ = 12.0, 3.2 Hz, 1H), 4.34 – 4.23 (m, 2H), 4.17 (dd, $J$ = 12.1, 6.8 Hz, 1H), 4.00 (t, $J$ = 6.1 Hz, 2H), 3.70 – 3.59 (m, 2H), 3.23 (s, 9H), 2.33 ($m_c$, 4H), 2.21 – 2.10 (m, 3H), 1.65-1.68 (m, 4H) 1.48 (dt, $J_{1,2}$ = 7.2 Hz, 2H), 1.44 – 1.22 (m, 40H), 1.18 – 1.04 (m, 4H), 0.90 (t, $J$ = 6.6 Hz, 3H) ppm.

$^{13}$C NMR{$^1$H} (101 MHz, $CD_3OD$) δ = 174.93, 174.43, 85.01, 71.83, 69.43, 67.49, 64.91, 63.62, 60.49, 54.69, 34.89, 33.84, 33.74, 33.09, 30.81, 30.78, 30.66, 30.49, 30.20, 29.98, 29.69, 29.64, 29.51, 26.02, 25.76, 25.74, 24.88, 24.67, 23.75, 19.00, 14.46 ppm.

$^{31}$P NMR{$^1$H} (162 MHz, $CD_3OD$) δ = -0.56 ppm.

HR-MS (ESI positive) m/z calculated for $C_{42}H_{78}N_3O_8P$: 783.5527; found: 784.558 $[M+H]^+$.

Yield: 13.4 mg (17.1 µmol, 18 %).

**Synthesis of PC(20:0/$Y_{16}$)**

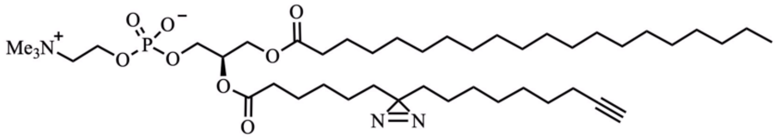

A solution of 31.0 mg of the bifunctional C-16 fatty acid ($Y_{16}$) (111µmol, 1.3 eq.) in 1 ml $CDCl_3$ was treated with 40.0 mg EDC·HCl (209µmol, 2.4 eq.) and 5.5 mg DMAP (45.0 µmol, 0.50 eq.). After 10 min, the solution of the activated fatty acid was added to a solution of 48.3 mg 1-*O*-icosanoyl-2-lyso-PC (87.4 µmol, 1.0 eq.) in 2 ml dry DMF and 1 ml $CDCl_3$. The reaction mixture was stirred overnight at room temperature. After removing the solvents under reduced pressure, the product was purified by flash chromatography ($CHCl_3$/MeOH/$H_2O$ 70:30:2). PC(20:0/$Y_{16}$) was isolated as a yellowish oil.

$^1$H NMR (400 MHz, $CD_3OD$) δ = 5.24 (s, 1H), 4.44 (dd, $J$ = 12.0, 3.2 Hz, 1H), 4.31-4.24 (m, 2H), 4.17 (dd, $J$ = 12.1, 6.8 Hz, 1H), 4.00 (t, $J$ = 6.0 Hz, 2H), 3.64 ($m_c$, 2H), 3.23 (s, 9H), 2.33 ($m_c$, 4H), 2.21 – 2.11 (m, 3H), 1.54 1.65-1.68 (m, 4H) 1.48 (dt, $J_{1,2}$ = 7.2 Hz, 2H), 1.44 – 1.22 (m, 44H), 1.13-1.08 (m, 4H), 0.90 (t, $J$ = 6.6 Hz, 3H) ppm.

$^{13}$C NMR{$^1$H} (101 MHz, $CD_3OD$) δ = 174.92, 174.43, 85.01, 71.94, 69.43, 67.45, 64.92, 63.62, 60.50, 54.68, 34.90, 33.84, 33.74, 33.09, 30.79, 30.49, 30.20, 29.98, 29.64, 26.02, 25.77, 24.88, 24.67, 23.75, 18.99, 14.45 ppm.*

*Signals between 29.5 and 30.8 ppm overlap and do not resolve individually.

$^{31}$P NMR{$^1$H} (162 MHz, $CD_3OD$) δ = -0.56 ppm.

HR-MS (ESI positive) m/z calculated for $C_{44}H_{82}N_3O_8P$: 811.5840; found: 812.590 $[M+H]^+$.
Yield: 5.1 mg (6.4 µmol, 7 %).

**Synthesis of 13-Oxohexadec-17-ynoic acid**

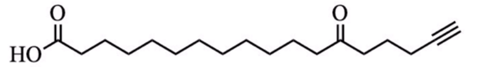

A solution of 5-Hexynoic acid (5.0 g, 44.6 mmol, 1.0 eq.) in 40 ml dry DCM was treated with 6 drops of dry DMF. Subsequently, oxalyl chloride (3.4 ml, 51mmol, 1.1 eq.) was added dropwise at 0 °C under an argon atmosphere. The reaction mixture was stirred for 20 min at 0 °C followed by 60 min at room temperature. A solution of 1-morpholinocyclododecene (15.0 g, 59.7 mmol, 1.3 eq.) in 20 ml dry DCM was heated to 35 °C in an open flask. After addition of $NEt_3$ (11 ml,78.9 mmol, 1.8 eq.) the acyl chloride mixture was added dropwise. The reaction mixture was stirred overnight at room temperature. 75 ml chloroform and aqueous HCl (50 ml, 20% v/v) were added and the mixture was stirred at room temperature for 4 h. The phases were separated and the organic phase was washed with water (3x). The combined aqueous layers were reextracted with DCM (3x) and the organic phases were then combined and their solvent removed under reduced pressure. The crude product was purified by flash chromatography (CyHex/EtOAc, 97:3). Product containing fractions were evaporated under reduced pressure and directly used in the next step. The crude intermediate (ca. 10 g) was heated to 100 °C and treated with aq. KOH (70 ml, 60 % v/v). After stirring for 15 min at 100 °C the reaction mixture was cooled to room temperature for 60 min. The reaction mixture was diluted with 200 ml water and slowly neutralised by dropwise addition of conc. aqueous HCl (20 ml). Upon neutralization, a colourless precipitate of crude product appeared which was collected by filtration. Further crude product was obtained by extracting the remaining water phase with chloroform (3x), combining the organic layers and removing the solvent under reduced pressure. The crude product was purified by flash chromatography (DCM/MeOH, 97:3 to 95:5). 13-Oxohexadec-17-ynoic acid was isolated as a white solid.
$^1$H NMR (400 MHz, $CDCl_3$) δ = 2.55 (t, $J$ = 7.3 Hz, 2H), 2.40 (t, $J$ = 7.5 Hz, 2H), 2.29 (t, $J$ = 7.5 Hz, 2H) 2.28 – 2.17 (m, 2H), 1.99 – 1.91 (m, 1H), 1.86 – 1.72 (m, 2H), 1.70 – 1.49 (m, 4H), 1.40 – 1.17 (m, 14H) ppm.
$^{13}$C NMR{$^1$H} (101 MHz, $CDCl_3$) δ = 210.91, 179.26, 83.80, 69.11, 43.16, 41.17, 34.02, 29.61, 29.51, 29.36, 29.33, 29.17, 24.82, 24.02, 22.40, 17.93 ppm. *
*Signals between 29.2 and 29.9 ppm overlap and do not resolve individually.
HR-MS (ESI negative) m/z calculated for $C_{18}H_{30}O_3$: 294.2195; found: 293.213 $[M-H]^-$.
Yield: 2.2 g (8.3 mmol, 18 % over 2 steps).

**Synthesis of bifunctional fatty acid $Y_{18}$**

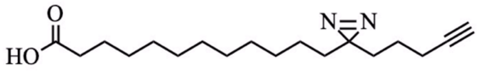

In a 250 ml flask molecular sieves (3Å, powder) were dried with heat under high vacuum. Ammonia (20 ml, 7N, solution in MeOH) was then added and a solution of 13-Oxohexadec-17-ynoic acid (2.0 g, 7.5 mmol, 1.0 eq.) in 25 ml dry MeOH was added dropwise. The reaction mixture was stirred at room temperature for 3 h. Subsequently, a solution of hydroxylamine-O-sulfonic acid (1.80 g, 15.9 mmol, 2.1 eq., dried under high vacuum for 3 h) was dissolved in 15 ml dry MeOH and the solution was added dropwise at 0 °C to the reaction mixture. The reaction mixture was stirred at room temperature overnight. Molecular sieves were removed by filtration and the filtrate was concentrated under reduced pressure. The residue was redissolved in 150 ml MeOH. 6 ml $NEt_3$ were added dropwise at 0 °C. Subsequently, iodine was added in small batches at 0 °C until the mixture remained brownish. The reaction was quenched with sat. aqueous $Na_2S_2O_3$-solution. The reaction mixture was extracted with DCM (3x), the combined organic layers were dried over $Na_2SO_4$ and the solvent was removed under reduced pressure. The crude product was purified by flash chromatography (DCM/MeOH 97:3). The product containing fraction were purified further by HPLC using a Macherey Nagel VP 250/32 Nucleodur C18 HTec column at 30 ml/min eluting with a gradient. The solvent system used was A (75 % MeOH, 25 %water, + 4 % AcOH) and B (50 % MeCN, 40 % i-propanol, 10 % MeOH, + 4 % AcOH). Gradient: 0-5 min: 0-45 % B, 5-15 min: 45-75 % B, 15-20 min: 75-100 % B, 20-27 min: 100-0% B, 27-30 min: 0 % B. The retention time of the product was found to be 17.2 min. The bifunctional fatty acid was isolated as a light yellowish oil.

$^{1}H$ NMR (400 MHz, $CDCl_3$) δ = 2.35 (t, *J* = 7.5 Hz, 2H), 2.16 (td, *J* = 7.0, 2.7 Hz, 2H), 1.97 – 1.91 (m, 1H), 1.71 – 1.58 (m, 2H), 1.55 – 1.44 (m, 2H), 1.42 – 1.15 (m, 18H), 1.15 – 0.99 (m, 2H) ppm.

$^{13}C$ NMR{$^{1}H$} (101 MHz, $CDCl_3$) δ = 178.14, 83.63, 69.00, 33.81, 33.02, 32.00, 29.63, 29.57, 29.51, 29.35, 29.19, 28.60, 24.83, 23.97, 22.91, 18.12 ppm.

HR-MS (ESI negative) m/z calculated for $C_{18}H_{30}N_2O_2$: 306.2307; found: 305.224 $[M-H]^-$.

Yield: 135.0 mg (440.0 µmol, 6 %)

#### Synthesis of GlcCer(Y)

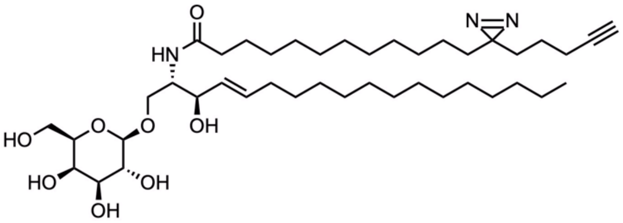

4.3 mg (14 µmol, 1.1 eq.) of the C18 bifunctional fatty acid was mixed with 5.9 mg (12.8 µmol, 1.0 eq.) of D-glucosyl-ß1-1'-D-erythro-sphingosine and 5.1 mg (33.3 µmol, 2.6 eq.) of 1-Hydroxybenzotriazole hydrate. The solids were dried over high vacuum and dissolved in 2 mL of dry DCM and 1 mL of dry THF. The reaction mixture was cooled to 0 °C and *N*-(3-Dimethylaminopropyl)-*N'*-ethylcarbodiimide (4.5 µl, 25.4 µmol, 2 eq.) was added. The reaction mixture was stirred under an argon atmosphere at room temperature for 48 h. The solvents were removed under reduced pressure and the crude product was purified with flash chromatography ($CHCl_3$/MeOH 20:1 to 10:1). The product GlcCer(Y) was isolated as a white solid.

$^{1}H$ NMR (400 MHz, $CD_3OD$) δ = 7.83 ($m_c$, 1H), 7.69 ($m_c$, 1H), 7.56 – 7.38 (m, 2H), 5.73 – 5.57 (m, 1H), 5.48 – 5.33 (m, 1H), 4.24 – 4.10 (m, 2H), 4.09 – 3.99 (m, 1H), 3.98 – 3.87 (m, 1H), 3.83 – 3.74 (m, 1H), 3.74 – 3.61 (m, 2H), 3.62 – 3.39 (m, 4H), 2.23 – 2.06 (m, 5H), 2.05 – 1.92 (m, 2H), 1.62 – 1.49 (m, 2H), 1.49 – 1.39 (m, 2H), 1.39 – 1.15 (m, 44H), 1.13 – 0.99 (m, 2H), 0.92 – 0.78 (m, 3H) ppm.

$^{13}C$ NMR{$^{1}H$} (101 MHz, $CD_3OD$) δ = 175.97, 134.97, 131.36, 129.57, 128.26, 127.12, 118.69, 111.45, 105.39, 84.12, 76.78, 74.86, 72.99, 72.65, 70.32, 70.05, 69.96, 62.53, 54.82, 37.38, 33.82, 33.46, 33.10, 32.79, 30.85, 30.81, 30.75, 30.66, 30.61, 30.55, 30.50, 30.43, 30.32, 29.21, 27.15, 24.91, 24.01, 23.76, 18.56, 14.46 ppm.

HR-MS (ESI positive) m/z calculated for $C_{42}H_{75}N_3O_8$:749.5554; found: 750.561 $[M+H]^+$.

Yield: 9.0 mg (12.0 µmol, 94 %).

### NMR Spectra

#### [1]H spectrum of PC(18:0|Y)

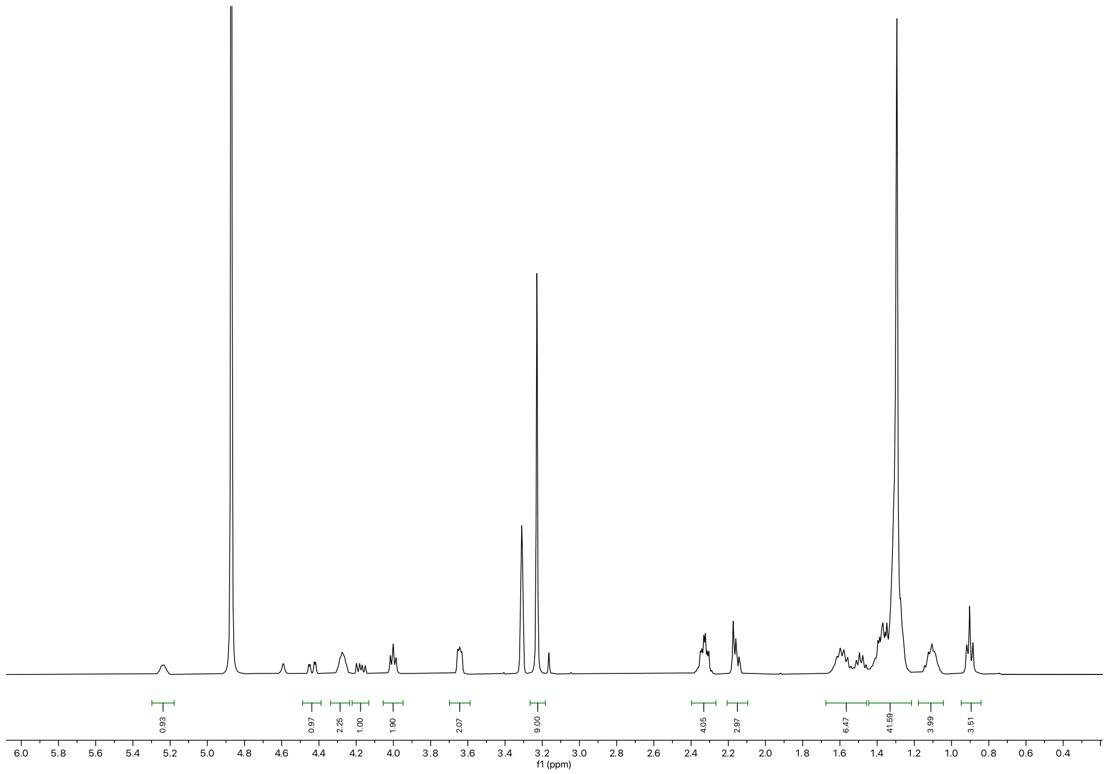

### $^{13}$C spectrum PC(18:0|Y)

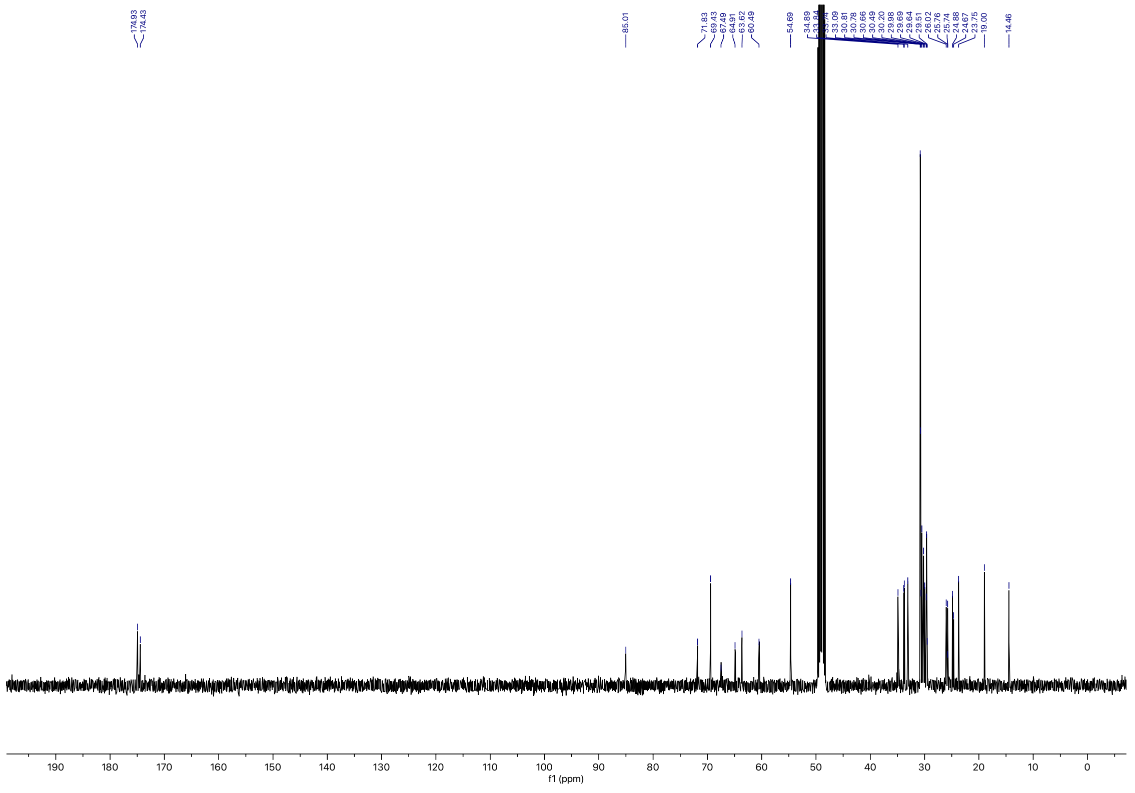

**$^{31}$P spectrum of PC(18:0|Y)**

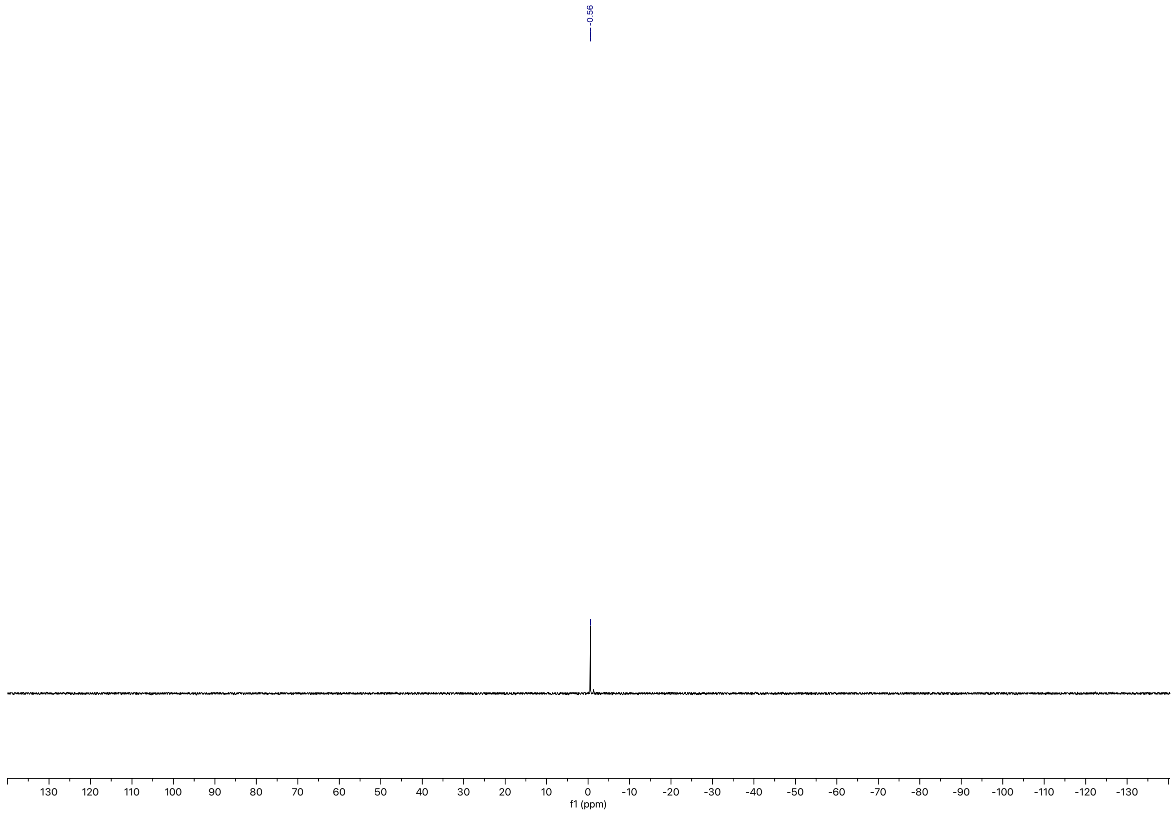

### [1]H spectrum of PC(20:0|Y)

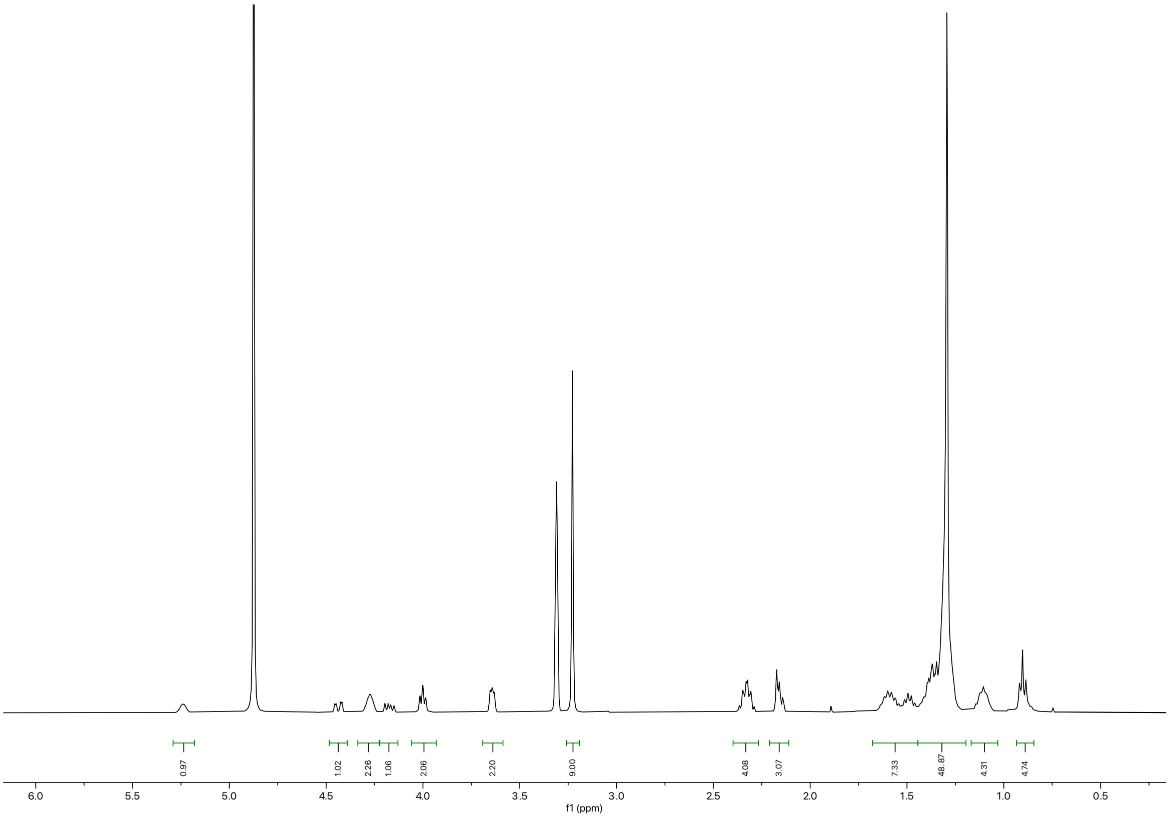

#### $^{13}$C spectrum of PC(20:0|Y)

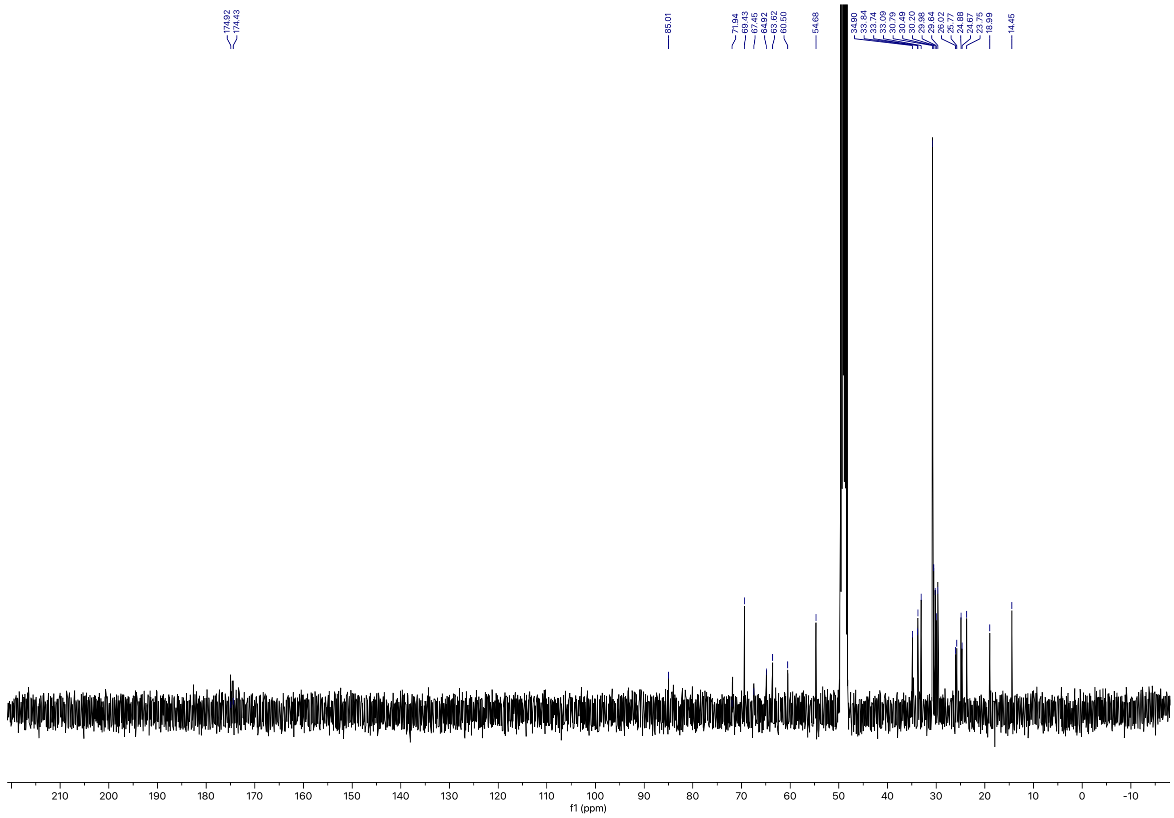

### $^{31}P$ spectrum of PC(20:0|Y)

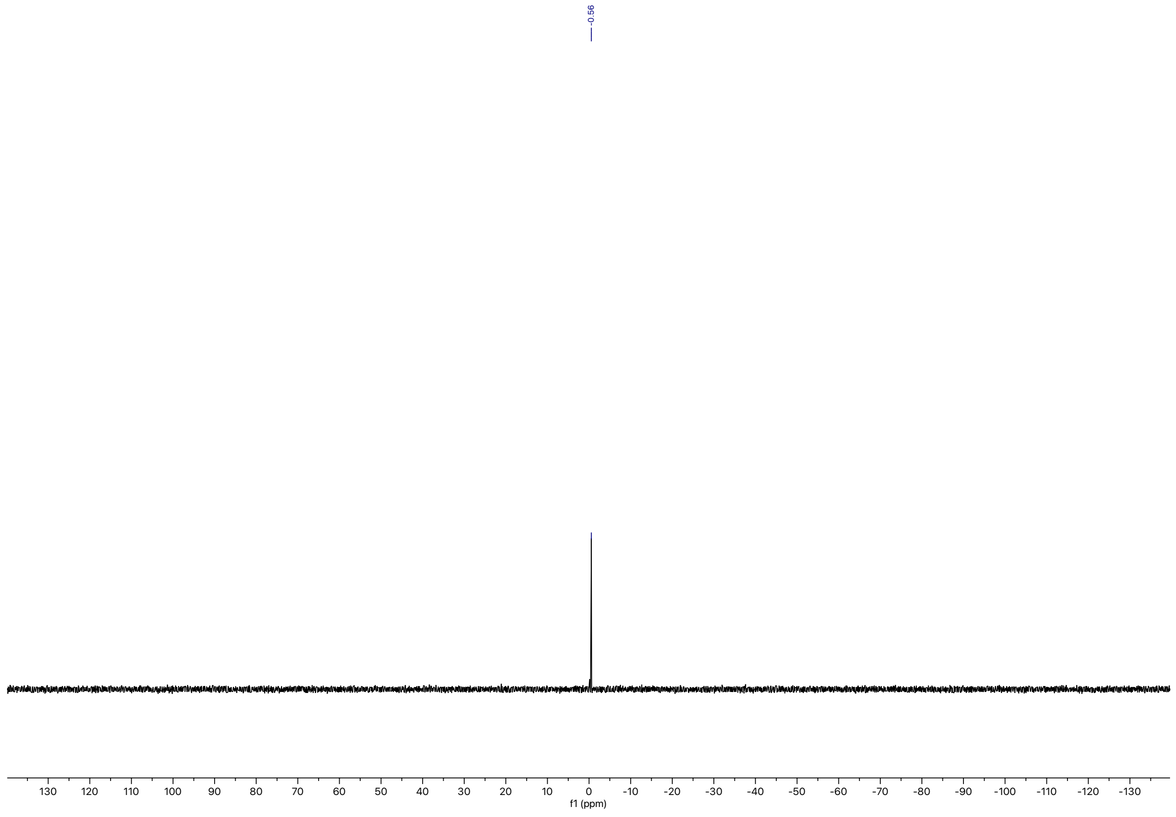

[1]H spectrum of 13-Oxooctadec-17-ynoic acid

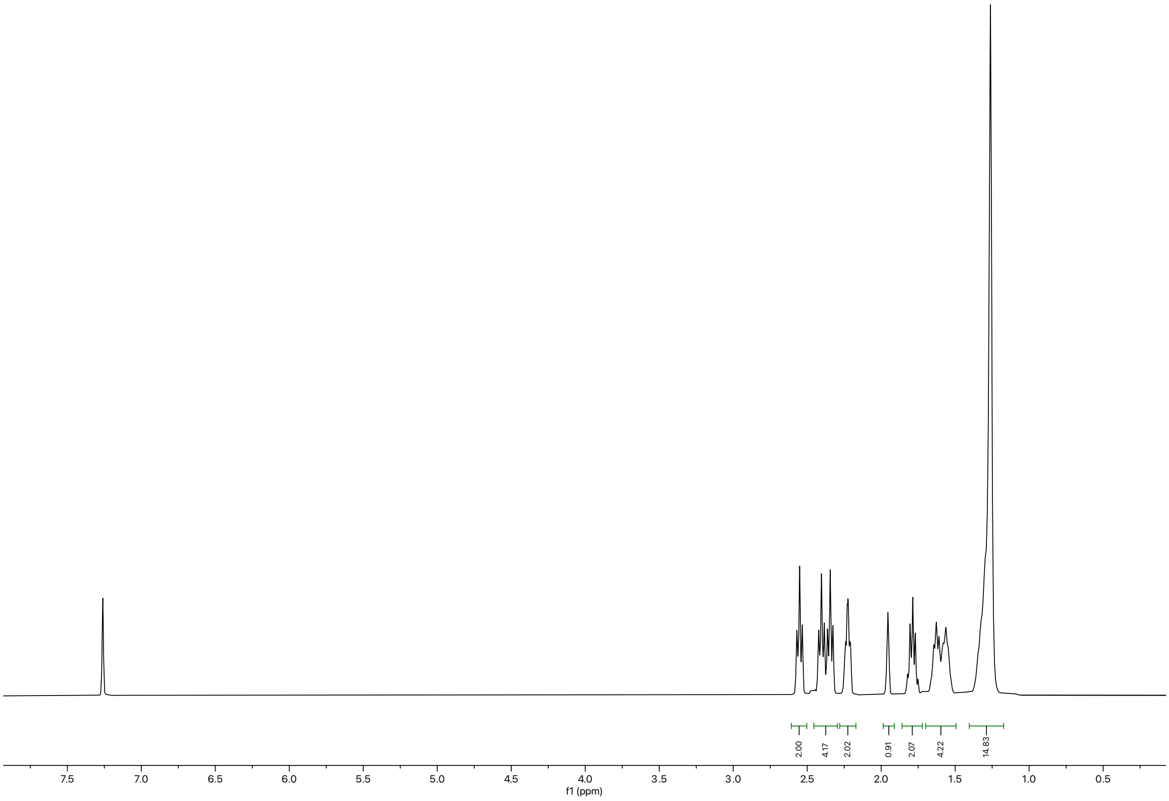

#### $^{13}$C spectrum of 13-Oxooctadec-17-ynoic acid

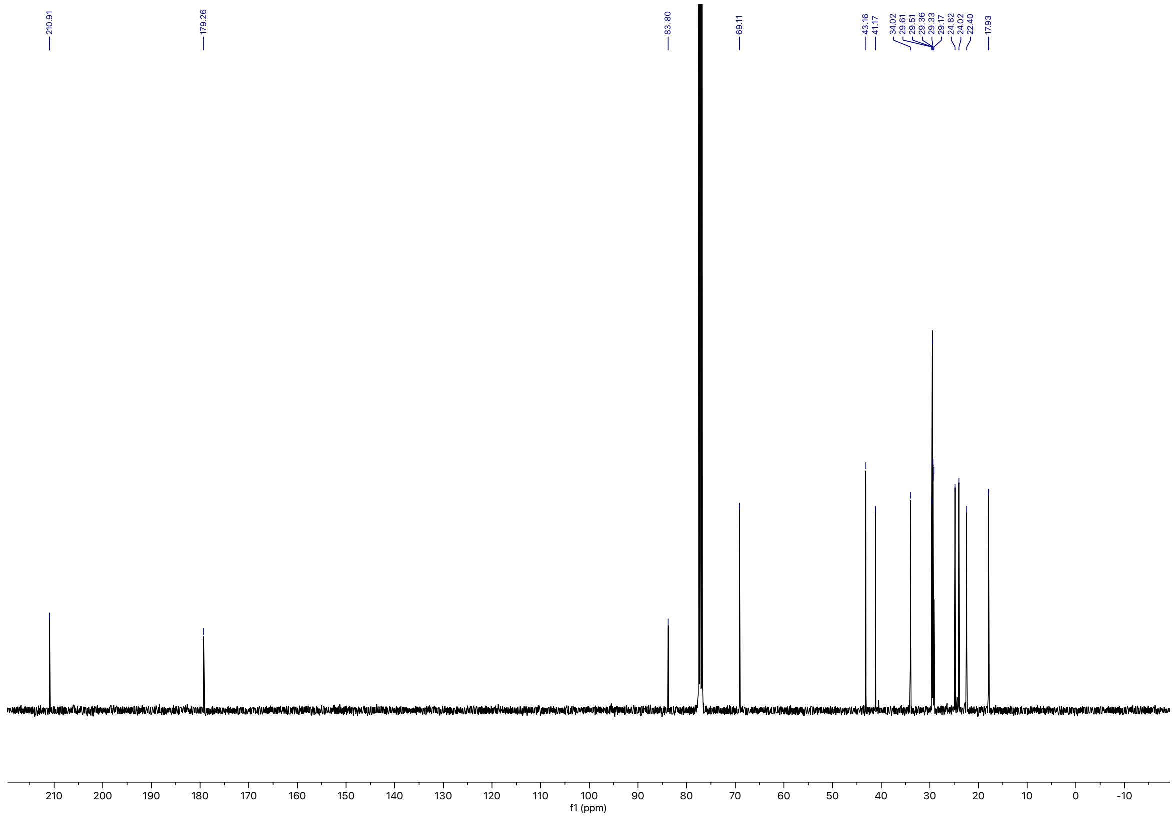

**$^{1}H$ spectrum of bifunctional fatty acid $Y_{18}$**

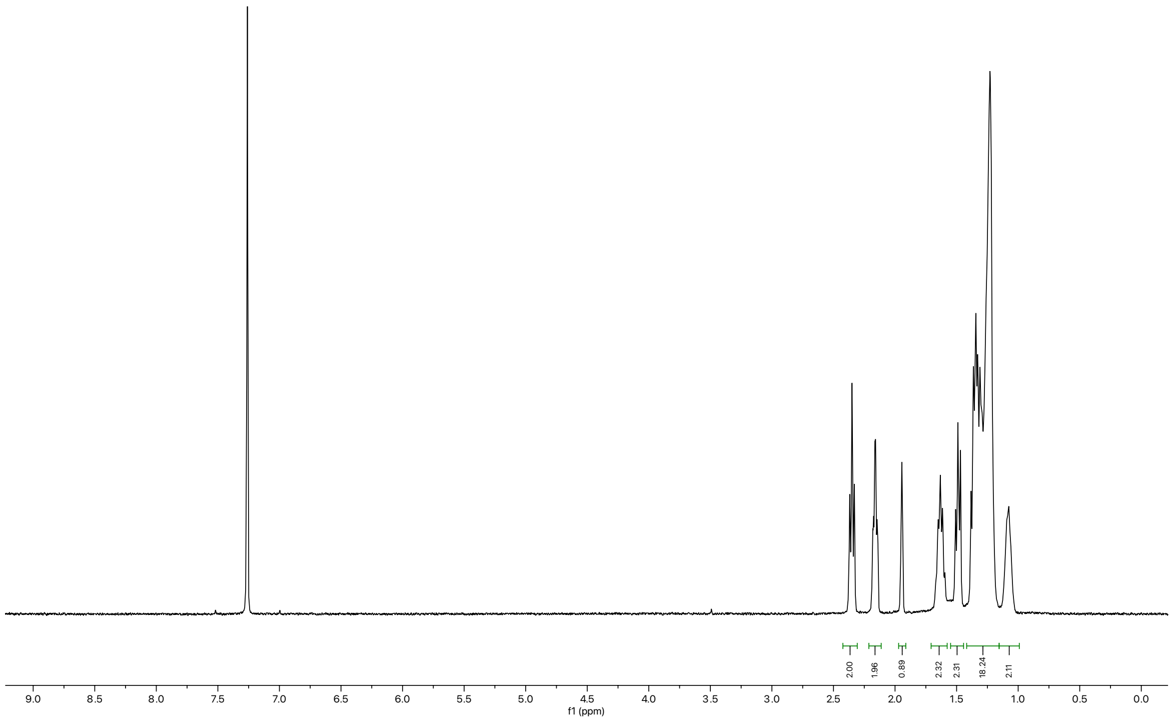

#### $^{13}C$ spectrum of bifunctional fatty acid $Y_{18}$

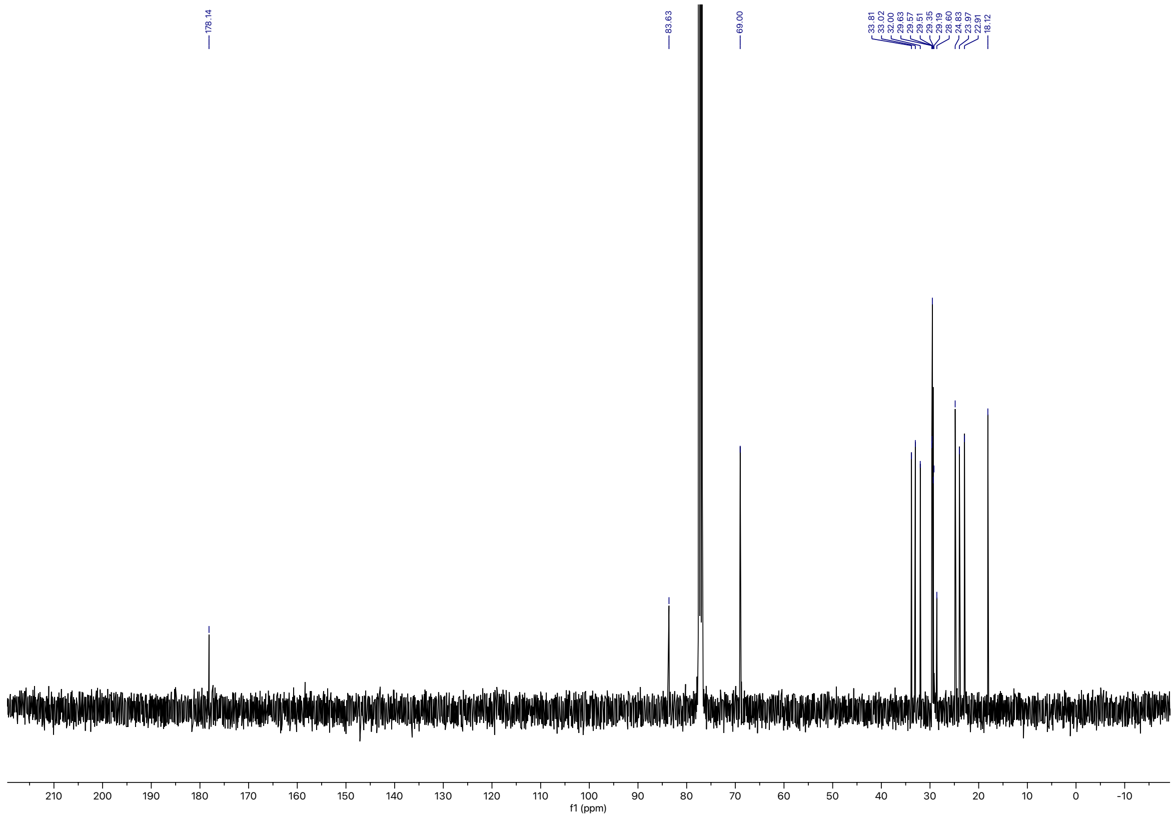

#### [1]H spectrum of GlcCer(Y)

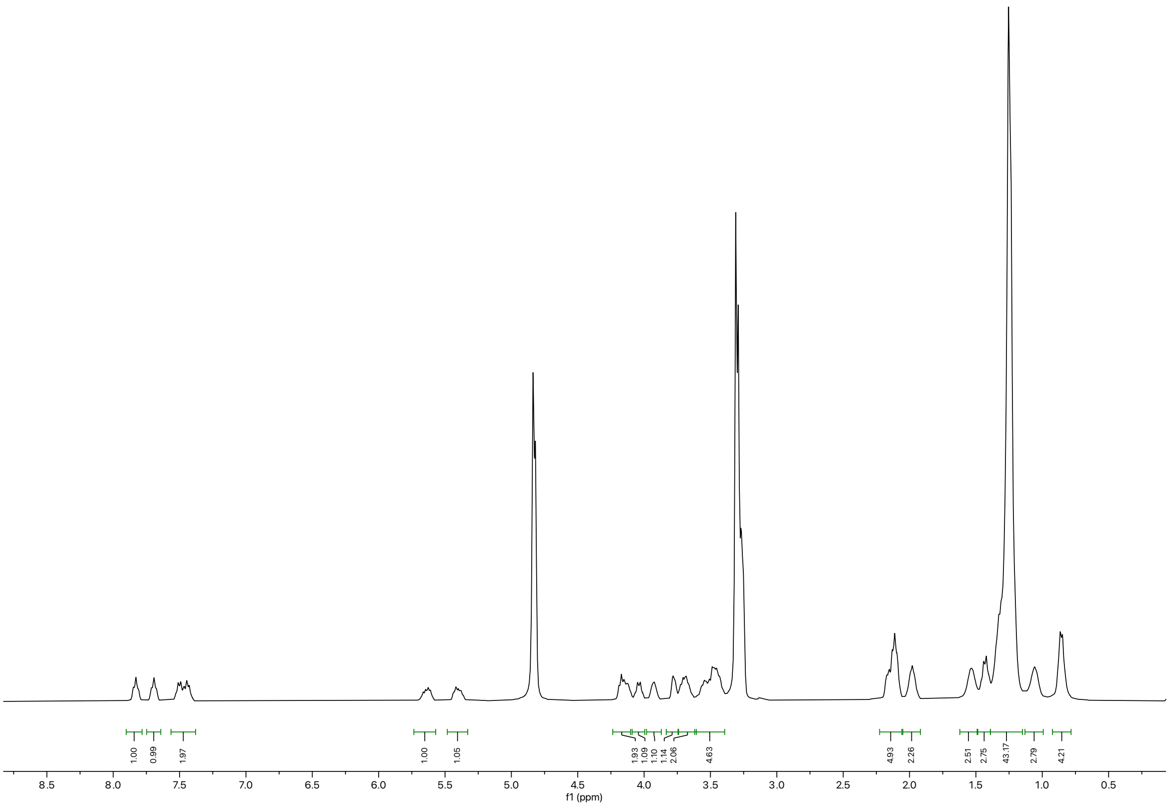

### $^{13}$C spectrum of GlcCer(Y)

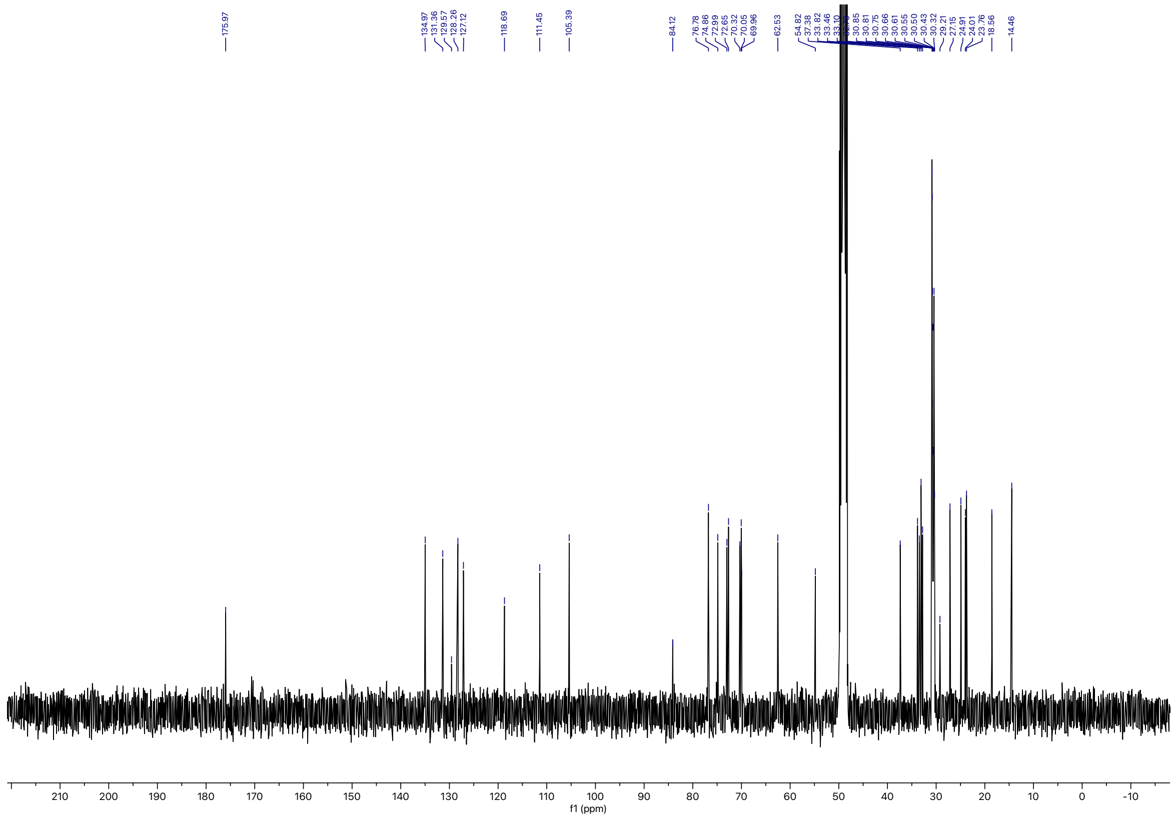

### Supplementary Figures and Data

#### Supplementary Data 1

**LED powers.** The power output of a 308nm LED as previously reported [1], and a high-power 365nm LED with and without collimating lens were measured over 10s. Values are reported in Table 2.

Table 2| Power measurements of different LEDs.

| LED type | Standard 308nm | High power 365nm (no lens) | High power 365nm(lens) |
|---|---|---|---|
| Power (mW) | 0.202 ± 6 | 1323 ± 11 | 1423 ± 15 |

A multi-LED strip suitable for synchronous photo-activation of 12 wells in a 96-well format was characterized for their power output per LED. The respective values are reported in Table 3.

Table 3| Power measurements of the multi-LED setup.

| LED ID | 1 | 2 | 3 | 4 | 5 | 6 | 7 | 8 | 9 | 10 | 11 | 12 |
|---|---|---|---|---|---|---|---|---|---|---|---|---|
| Power (mW) | 2056 ± 6 | 1950 ± 10 | 1896 ± 6 | 1850 ± 1 | 1967 ± 6 | 1897 ± 6 | 1910 ± 0 | 1987 ± 6 | 2057 ± 6 | 2040 ± 10 | 2047 ± 6 | 1973 ± 20 |

#### Supplementary Data 2: An estimation for geometry-based sorting of lipids into curved membranes

The Helfrich membrane bending energy [11] $E = \frac{1}{2}kA(C - Co)^2$ describes the energy cost to bend a surface area A of a membrane from its resting curvature $C_0$ is dependent on both the stiffness k of the membrane as well as the final curvature C. Both $C_0$, k, and the total bending energy depend on the lipid composition. In order to reduce the bending energy, lipids can either rearrange in order for $C_0$ to match C or to locally reduce k [12–16]. For determining the sorting of lipids with $C_0$ from a flat membrane to a bent membrane we calculate: $\Delta E = E_{flat} - E_{curved} = \frac{1}{2}ka[(C_{flat} - C_0)^2 - (C_{curved} - C_0)^2]$, with a boing the surface area of the lipid. We estimate the following values: k = ~20 $k_BT$; a = ~0.5 $nm^2$, $C_{flat}$ = 0 nm-1; $C_{curved}$ = 1/50 $nm^{-1}$ or 1/25 $nm^{-1}$. Results are summarized in Table 4.

Table 4| Sorting energies of lipids into bend membranes.

| **lipid** | **$C_0$ [$nm^{-1}$]** | **$\Delta E$ [$kT$] for $C_{curved}$ 1/25 $nm^{-1}$** | **$\Delta E$ [$kT$] for $C_{curved}$ 1/50 $nm^{-1}$** | **ref** |
|---|---|---|---|---|
| PE 16:0-18:1 | -0.0317 | -0.0207 | -0.0083 | [17] |
| PC 16:0 16:0 | 0.0050 | -0.0060 | -0.0010 | [17] |
| PC 16:0 18:1 | 0.0010 | -0.0076 | -0.0018 | [17] |
| SM | 0.0000 [e] | -0.0080 | -0.0020 | |

[e] – value was estimated based on similar lipids or literature

#### Supplementary Data 3: Values used for melting temperatures $T_m$ of lipid species

The melting temperatures ($T_m$) of all experimentally assessed lipid species were determined from the literature, if possible. Values used in this study are summarized in Table 5.

Table 5| Melting temperatures of selected lipid species

| lipid | $T_m$ [°C] | ref |
|---|---|---|
| GlcCer | 80 | [18] |
| SM | 41 | [19] |
| PC 16:0-16:0 | 42 | [20,21] |
| PC 16:0-18:0 | 49 | [20,21] |
| PC 16:0-20:0 | 57[e] | |
| PC 16:0-18:1 | -2 | [20,21] |
| PC 16:0-20:4 | -22.5[e] | [21] |
| PC 20:4-16:0 | -22.5 | [21] |
| PE 16:0-18:1 | 25 | [20] |
| pPC 16:0-18:1 | / | |

[e] – value was estimated based on similar lipids

#### Supplementary Data 4: An estimation for asymmetry-based sorting of lipids into curved membranes

We assume that lipid asymmetry is maintained in CCPs and that lipid probes have reached their steady state trans-bilayer distribution at the time point of photocrosslinking. We used the mol% of distinct lipid species per total phospholipids on the cytosolic leaflet and extracellular leaflet from Lorent et al. [22], which are shown in Table 6.

Table 6| Asymmetric lipid leaflet distributions across the plasma membrane for selected lipid species

| lipid | Cyto [mol%] | Extra [mol%] | ref |
|---|---|---|---|
| GlcCer | / | / | |
| SM | 0.060 | 1.203 | [22] |
| PC 16:0-16:0 | 1.301 | 0.466 | [22] |
| PC 16:0-18:0 | 1.004 | 0.000 | [22] |
| PC 16:0-20:0 | 1.000[e] | 0.000[e] | [22] |
| PC 16:0-18:1 | 4.716 | 6.353 | [22] |
| PC 16:0-20:4 | 0.982[e] | 3.589[e] | [22] |
| PC 20:4-16:0 | 0.982 | 3.589 | [22] |
| PE 16:0-18:1 | 3.111 | 0.135 | [22] |
| pPC 16:0-18:1 | / | / | |

Next, we estimated the phospholipid content of the cytosolic and extracellular leaflet of a vesicle with an outer diameter (cytosolic, short: cyto) of 100 nm. The vesicle functions as a proxy for the highly curved late-stage CCP just before fission. We assume a bilayer thickness of 4 nm, and thus an inner (extracellular, short: extra) diameter of 92 nm. Thus, we calculate the surface areas $A_{cyto} = 31416$ nm$^2$ and $A_{extra} = 26590$ nm$^2$. We assume that 50% of the surface area (15708 nm$^2$) is covered by proteins of cylindrical shape. The surface area per phospholipid is estimated to 0.5 nm$^2$. Finally, we consider the asymmetric distribution of cholesterol, which drives an asymmetric distribution of phospholipids across both leaflets with an estimate of 42 mol% of all plasma membrane lipids on the cytosolic side and 22 mol% of all plasma membrane lipids on the extracellular side. These values were taken as estimates from Doktorova et al. [23]. Taken together, this renders a total of 13020 phospholipids to the cytosolic side of the vesicle and 4840 to the extracellular side. We similarly estimate lipid numbers for a flat membrane with the surface area of $A_{flat} = 31416$ nm$^2$ to $\#_{cyto} = 13020$ and $\#_{extra} = 6820$.

To determine the numbers of lipid species per leaflet for a flat membrane and a curved membrane we take the product of the total estimated phospholipid number per leaflet with the previously reported mol% for the respective lipid species and leaflet as shown Table 7. Finally, the total estimated number is evaluated for the flat membrane and curved membrane (pit) and their ratio of the lipid number in the curved membrane over the number of lipids in the flat membrane (Ratio) is taken similar to the analysis to determine lipid pit enrichments for the STED data.

Table 7| Ratio of lipid content in a bend to a flat membrane.

| | **Flat membrane** | | | **Curved membrane** | | | |
|---|---|---|---|---|---|---|---|
| **lipid** | **#cyto** | **#extra** | **#total** | **#cyto** | **#extra** | **#total** | **Ratio** |
| SM | 7.81 | 82.04 | 89.86 | 7.81 | 58.23 | 66.04 | 0.73 |
| PC 16:0-16:0 | 169.39 | 31.78 | 201.17 | 169.39 | 22.55 | 191.94 | 0.95 |
| PC 16:0-18:0 | 130.72 | 0.00 | 130.7208 | 130.72 | 0.00 | 130.72 | 1.00 |
| PC 16:0-20:0 | 130.20 | 0.00 | 130.2 | 130.20 | 0.00 | 130.20 | 1.00 |
| PC 16:0-18:1 | 614.02 | 433.27 | 1047.30 | 614.02 | 307.49 | 921.51 | 0.88 |
| PC 16:0-20:4 | 127.86 | 244.77 | 372.62 | 127.86 | 173.71 | 301.56 | 0.81 |
| PC 20:4-16:0 | 127.86 | 244.77 | 372.62 | 127.86 | 173.71 | 301.56 | 0.81 |
| PE 16:0-18:1 | 405.05 | 9.21 | 414.26 | 405.05 | 6.53 | 411.59 | 0.99 |

#### Supplementary Figure 1

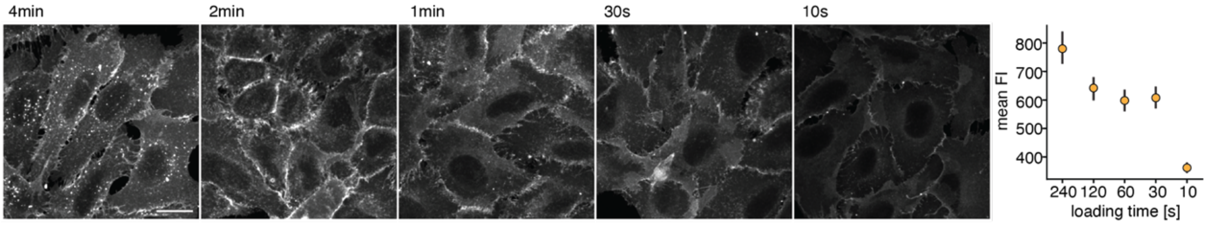


**Different lipid loading times**. Optimization of lipid loading (pulse) times for high temporal resolution of lipid uptake into early endosomes. U2OS wildtype cells were loaded with bifunctional sphingomyelin for 4min, 2min, 1min, 30s and 10s. Mean fluorescent intensities are plotted with the respective standard deviations. Scale bar in 20 µm.

#### Supplementary Figure 2

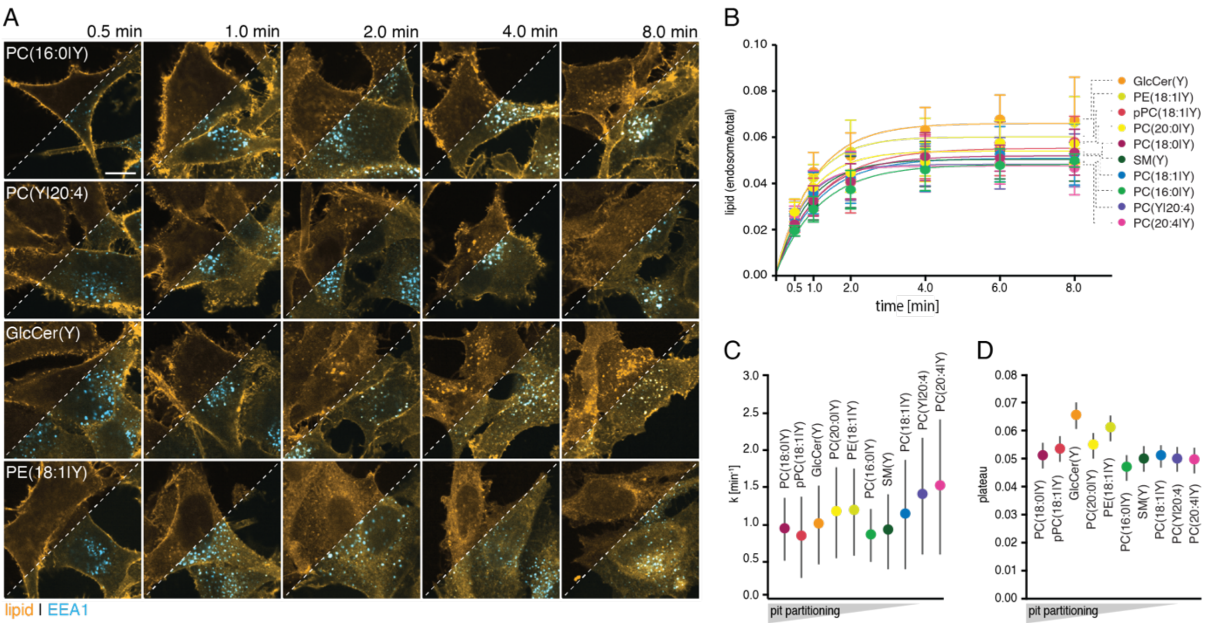


**Lipid uptake into early endosomes reveals moderate sorting during endocytosis.** A: Exemplary fluorescent images of SK-MEL2 wildtypes cells loaded with lipid for 0.5min and chased for 0min, 0.5min, 1.5min, 3.5 min, and 7.5min are shown. The left half of each image shows the lipid channel only (orange), and the right half the overlay with the EEA1 co-stain (blue). Scale bar: 10µm. B: Relative fluorescent lipid signal in early endosomes over total lipid signal was quantified for six different time points and 10 lipids for 3 biological replicates. Mean and standard deviation are shown. Uptake traces were fitted to a one-phase association model. The order of lipids from highest to lowest early endosomal amount at 8min is shown. C: The rate constants determined from the fit are plotted with the 95% confidence interval shown. Rate constants are ordered from highest to lowest pit partitioning. D: The amplitudes of each fit are shown with the 95% confidence interval indicated. Amplitude values are ordered from highest to lowest pit partitioning.
